# Isoindole-benzohydrazide derivative potentiates colistin activity to overcome resistance conferred by MCR-1 variants in *Escherichia coli*

**DOI:** 10.64898/2026.10.10.758198

**Authors:** Anna S. Wolf, Ruth O. Ndupu, John N. Alumasa

## Abstract

Polymyxin antibiotic resistance mediated by the mobilized colistin resistance (MCR) enzyme threatens the last-resort treatment of deadly multidrug-resistant pathogens, including *Acinetobacter baumannii,* and calls for urgent mitigation efforts. We used a multifaceted approach involving computational and wet-lab techniques to identify an isoindole-benzohydrazide derivative, JNAL-042, which potentiates colistin activity in an *Escherichia coli* MCR-resistance phenotype screening model. JNAL-042 lacked intrinsic antibiotic activity but potentiated that of polymyxin antibiotics at 12-24 µg/ml, as demonstrated by bacterial growth susceptibility screens. Cell viability counts and time-kill assays corroborated these findings, demonstrating complete abolition of MCR-mediated resistance by the antibiotic-adjuvant combination at concentrations mirroring those of empty vector controls. Post-induction treatment with this combination similarly prevented the propagation of the resistance phenotype, albeit at a higher antibiotic dose. This landmark study presents JNAL-042 as a promising antibiotic adjuvant to combat MCR-mediated resistance and offers a viable solution to preserve last-resort polymyxin antibiotics.

**Graphical abstract:** 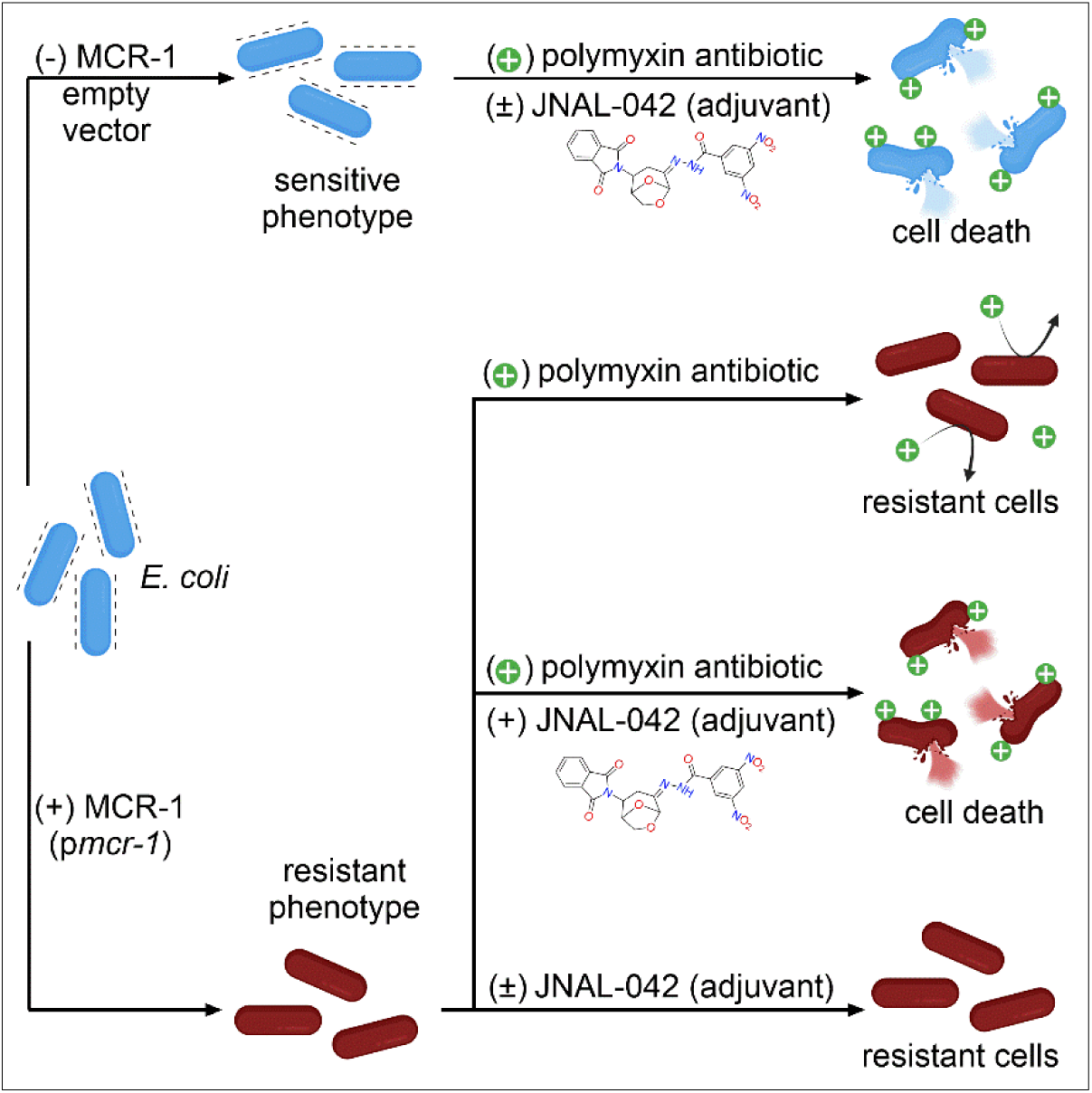

**Highlights:**

- Computational screen identified colistin adjuvant against MCR-1-mediated resistance
- Colistin-adjuvant combination abolished MCR-mediated resistance in time-kill assays
- Adjuvant rescued colistin activity in delayed-treatment kinetic time-kill assays
- Adjuvant lacked intrinsic activity and restored polymyxin activity in WT *E. coli*

## INTRODUCTION

Antibiotic-resistant bacteria remain a serious threat to human health, especially given the emergence of new resistance mechanisms, persistence of existing ones, and a depletion of the stockpile of effective antibiotics.^1,2^ Over one million fatalities occur annually from such infections globally, and this number is projected to increase drastically if left unresolved, with about 40 million deaths predicted by 2050.^3^ These alarming statistics call for additional effort to develop tools to combat this phenomenon. Nonessential enzymes are among the main arsenal of weaponry bacteria use to confer resistance to major antibiotic classes used in clinical settings.^4–6^ Genes encoding these enzymes are typically located on mobile genetic elements such as resistance plasmids, which facilitate easy cross-species dissemination.^7^ Enzyme-mediated resistance manifests primarily through structural modification of antibiotics or their molecular targets within bacterial cells.^5,8,9^ Studies on select classes of these enzymes have used antibiotic adjuvants to curb their resistance-conferring effects.^10^ Target-modifying enzymes mostly affect bacterial membranes or ribosomal structure, strategically altering the chemical composition of nucleic acids, lipids, or proteins near an antibiotic’s binding site(s), thereby conferring resistance.^6^ Membranes are the first line of defense for bacteria against harmful molecules and environmental stressors. Therefore, bacteria will go to extreme lengths to prevent damage when membrane integrity is compromised.^11^ Resistance to polymyxin antibiotics, including colistin (COL) and polymyxin B (PMB), which is in part mediated by the mobilized COL resistance enzyme (MCR), provides a chilling example of such a response.^12–14^

COL is a membrane-targeting cationic lipopeptide and a last-resort antibiotic, used to treat extremely severe infections caused by multidrug-resistant pathogens such as *Klebsiella pneumoniae* and *Acinetobacter baumannii*.^15–17^ This antibiotic acts by electrostatically binding to the negatively charged lipopolysaccharides (LPS), an essential component of the outer membrane (OM) of Gram-negative bacteria.^18^ This interaction displaces associated cations (e.g., Ca^2+^), destabilizing the OM, leading to membrane lysis and ultimately cell death.^15^ Plasmid-mediated resistance to COL is attributed to the translation product of the recently identified *mcr-1* gene.^12,19,20^ This gene encodes a membrane-bound zinc-metalloenzyme, MCR-1, with an integral N-terminal domain anchored in the inner leaflet of the cell wall. This section is connected via a linker to a catalytic C-terminal domain localized within the periplasm of Gram-negative bacteria.^21^ This enzyme obtains phosphoethanolamine (PEtN), primarily from phosphatidylethanolamine (PE), and transfers it to a phosphate group on lipid A, an essential component of LPS.^19,21^ PEtN transfer covalently modifies LPS, structurally altering the OM by introducing a positively charged amine.^21^ This structural change neutralizes the negatively charged membrane, preventing COL binding and conferring resistance.^21^ The *mcr* gene is evolving and spreading rapidly, with several variants displaying varying levels of polymyxin resistance being reported in pathogens such as *Acinetobacter baumannii and Pseudomonas aeruginosa.*^22^ Given that these genes are localized on highly mobile resistance plasmids, these trends call for new strategies to tackle this issue and restore the activity of polymyxins, a vital class of last-resort antibiotics.

Therapeutic targeting of nonessential resistance-conferring enzymes is a resourceful approach that employs adjuvants to potentiate the activity of affected antibiotics.^23,24^ Documented neutralization of β-lactamase-mediated hydrolysis of β-lactam antibiotics using adjuvants validates this approach.^25^ A few reports highlight the therapeutic targeting of MCR-1, including the use of metals such as silver, natural products, or combinations of polymyxins with existing FDA-approved drugs.^26–30^ Some of these studies show promising outcomes, but with caveats that make the feasibility of clinical application uncertain. For instance, studies combining known antibiotics or other drugs with COL have identified promising candidates for combinatorial therapy.^26,30^ Yet, a variant of MCR-1 has already displayed co-resistance to β-lactam antibiotics, and some of the drugs may enhance the established cytotoxic effects of polymyxins.^30,31^ While the approach is valid, its long-term implementation may be counterproductive, and the associated risks of cross-resistance or enhanced toxicity warrant alternative approaches. The use of computer-aided drug discovery (CADD) techniques to identify therapeutic agents is a versatile strategy with proven success across multiple bacterial target-based screening efforts.^32,33^ Contrary to traditional high-throughput screening methods, this technique is fast, cost-effective, and has a moderately high success rate in identifying effective therapeutics.^32^ Moreover, the availability of virtual small-molecule libraries with highly diverse chemical scaffolds, freely accessible to researchers, further justifies the use of CADD for target-based discovery efforts.^34^ Such libraries contain a treasure trove of novel bioactive molecules with therapeutic potential.

Here, we combined CADD with wet-lab methods to validate MCR-1 variants as viable drug targets for developing adjuvant-based therapy to combat MCR-mediated polymyxin resistance in Gram-negative bacteria. Using CADD, we identified an isoindole-benzohydrazide derivative, JNAL-042, that potentiates COL activity to abolish MCR-1-mediated resistance in efflux-deficient and wild-type *E. coli* cells. JNAL-042 was not cytotoxic to human embryonic kidney cells and showed no intrinsic antibiotic activity at any evaluated concentration when tested alone, reducing the likelihood of resistance developing over time. This groundbreaking study characterizes the biological activity of JNAL-042, demonstrating its potential for use in combination with COL to develop antibiotic adjuvant-based therapy against MCR-mediated resistance in Gram-negative bacterial pathogens.

## RESULTS

### Strain construction and antibiotic susceptibility assays establish *E. coli* phenotypic screening platform

The rapid evolution of the *mcr-1* gene and its implications for treating multidrug-resistant pathogens that rely on polymyxins as last-resort antibiotics prompted this study, which sought to validate MCR variants as viable drug targets for adjuvant-based therapy. We initiated this study by developing *E. coli* working models carrying the genes for two MCR variants, the wild-type (MCR-1) and a double mutant isoform (M6–P188A, P195S), and an empty vector control (VC). We primarily investigated this MCR-M6 variant based on reports that it confers dual resistance to polymyxins and β-lactam antibiotics.^31^ Strains were developed by transforming cloned plasmids containing the corresponding genes into the WT BW25113 or an efflux-deficient version of this strain (BW25113 Δ*tolC*::*cat*^35^). We used this latter option to maximize our success rate for identifying potential MCR-1 inhibitors by avoiding efflux pump effects, which could generate false-negative results. We validated the phenotypic traits and functionality of these strains using antibiotic susceptibility screens to compare dose-response and minimum inhibitory concentrations (MICs) of polymyxin, β-lactam, and lincosamide antibiotics. An efflux-deficient strain expressing the MCR-M6 variant showed a 3- to 10-fold increase in COL and PMB MICs relative to controls (**Figure 1A, D**). These values increased dramatically in WT strains, resulting in a 20–30-fold MIC increase relative to a control strain (**Table 1**). We observed a modest 2-fold MIC increase over the control strain for two β-lactam antibiotics, ampicillin (AMP) and cefoxitin (CXT) (**Figure 1B, D**), confirming this previously reported phenomenon.^31^ Importantly, the expression of this enzyme did not adversely affect the activity of clindamycin, a lincosamide antibiotic that targets protein synthesis, validating the selectivity of MCR-M6-mediated resistance **(Figure 1C, D)**. These results established our phenotypic screening platform, paving the way for identifying polymyxin potentiators against MCR-mediated resistance.

**Figure 1.**
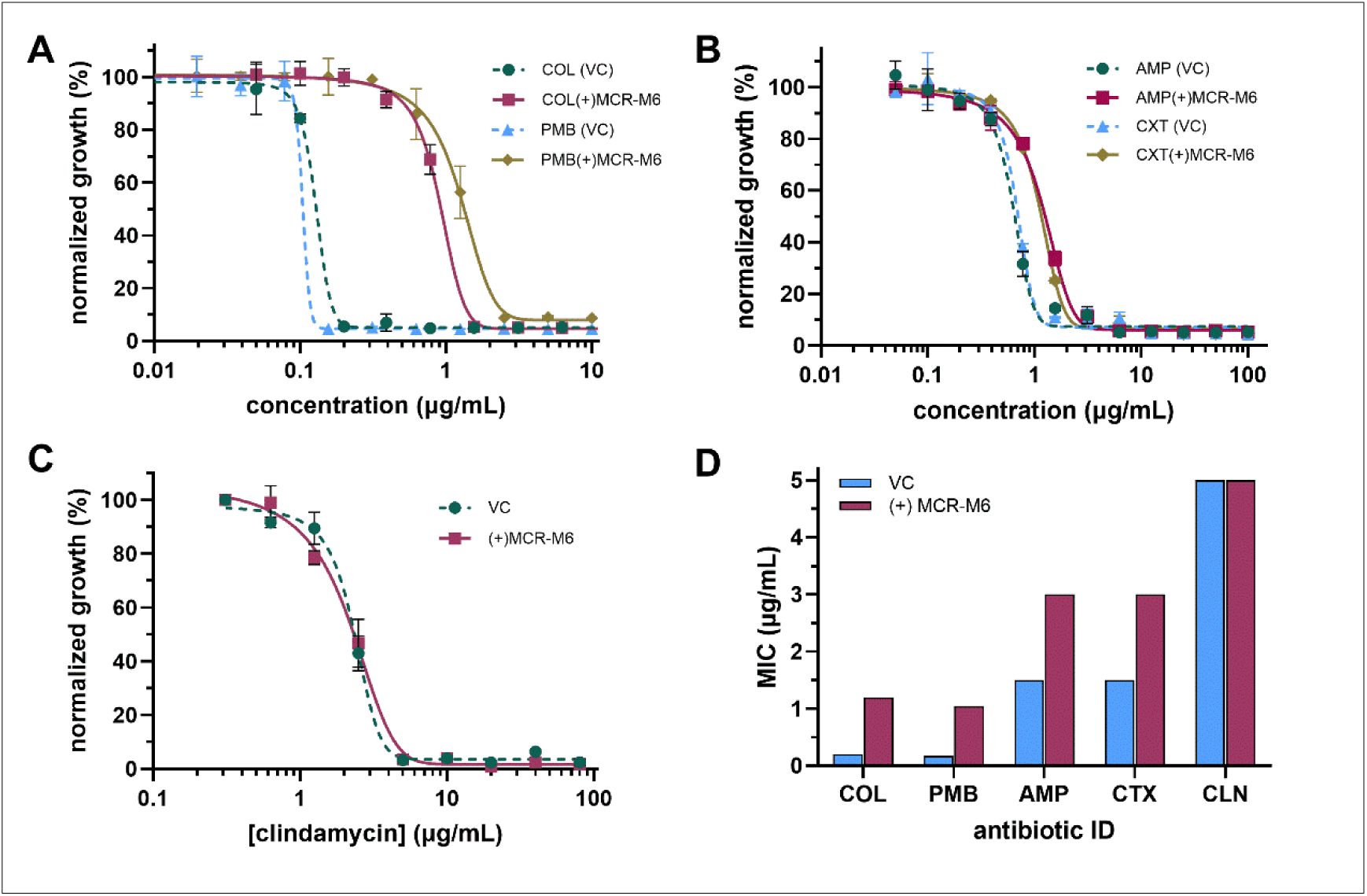
Antibiotic susceptibility screens confirm selectivity of MCR-M6-mediated resistance toward polymyxin and β-lactam antibiotics, but not lincosamides. The experiments utilized strains ASW-018 and ASW-021 carrying the *mcr-M6* gene or an empty vector control (VC) plasmid, respectively. A) Dose-response plots for polymyxin antibiotics, COL and PMB. B) Dose-response plot for β-lactam antibiotics, ampicillin (AMP) and cefoxitin (CXT). C) Dose-response plot for a lincosamide antibiotic, clindamycin (CLN). D) Average representative antibiotic MICs.

**Table 1.** Minimum inhibitory concentrations (MICs) for COL and PMB with or without JNAL-042 (12 µg/mL or 24 µg/mL for the efflux-deficient or WT strains, respectively).

| STRAIN ID | STRAIN CODE | PLASMID | COLISTIN MIC<sup>s</sup> (µg/mL) |  | POLYMYXIN B MIC<sup>s</sup> (µg/mL) |  |
| --- | --- | --- | --- | --- | --- | --- |
|  |  |  | (-) JNAL-042 | (+) JNAL-042 | (-) JNAL-042 | (+) JNAL-042 |
| WT *E. coli* | ASW-022 | pVC | 0.22 (0.08) | ND | 0.1 (0.05) | ND |
| WT *E. coli* | RON-011 | pMCR-1 | 1.25 (0) | 0.63 (0) | 1.25 (0) | 0.78 (0.31) |
| WT *E. coli* | ASW-020 | pMCR-M6 | 3.33 (0) | 1.56 (0.21) | 2.12 (0) | 0.44 (0.15) |
| *E. coli* ΔtolC::cat | ASW-021 | pVC | 0.17 (0.03) | ND | 0.07 (0.01) | ND |
| *E. coli* ΔtolC::cat | RON-001 | pMCR-1 | 0.33 (0) | 0.1 (0) | 0.26 (0.01) | 0.09 (0.04) |
| *E. coli* ΔtolC::cat | ASW-018 | pMCR-M6 | 1.19 (0.27) | 0.19 (0.03) | 1.04 (0.03) | 0.10 (0.05) |

<sup>s</sup>n ≥ 3; ND: not determined

### Virtual natural product-library screen identifies candidate compounds with adjuvant activity against MCR-1-mediated resistance

Motivated by our recent success in utilizing CADD to identify adjuvants against a resistance-conferring methyltransferase,^36^ we employed the same strategy to screen for potential inhibitors of MCR-1 activity. We conducted in silico screens to identify small molecules predicted to bind to the active site or substrate-binding regions of MCR-1.^37^ We focused on a virtual natural product library from the ZINC20 database, containing a subset of 10,191 compounds,^38^ hypothesizing that it would include unique, highly complex compounds, as typically observed in FDA-approved drugs from natural sources. We then screened compounds using AutoDock Vina with a cryo-EM structure of MCR-1 (PDB ID: 9NWW) in a substrate-bound conformation.^37^ These screens identified 1,682 compounds of interest based on predicted binding energies with a cutoff at -7.5 kcal/mol. These compounds were subsequently subjected to further in silico assessment using the Swiss-ADME online platform to evaluate predicted pharmacokinetic and drug-likeness properties.^39^ Ultimately, we purchased 12 top-ranked compounds based on their combined high binding affinity, drug-likeness, and structural diversity (**Figure S1**). This selection also prioritized predicted binding at the PE or LPS binding sites (**Figure 2A**), suggesting potential competitive binding with one of the MCR-1 substrates. We initially evaluated these compounds for intrinsic antibiotic activity against ASW-021, a TolC mutant strain carrying the empty vector. A 2-dose growth susceptibility assay at 50 or 100 μM identified four compounds with this property, suggesting they target essential bacterial components (**Figure 2B**). We excluded these four from the study and evaluated the remaining compounds with or without COL in a strain expressing the MCR-M6 variant to assess synergy. We added COL to one test plate at 1 µg/mL, a concentration that was typically ineffective against a bacterial strain expressing this variant. These experiments were crucial for corroborating the lack of intrinsic antibiotic activity in most remaining test compounds and identifying a false positive from the initial screen (compound 10) (**Figure 2C**). We eliminated this compound and others that lacked or showed very mild COL-potentiating properties from the study (**Figure 2C, D**). Intriguingly, our strategy identified a promising candidate, compound 3 (henceforth named JNAL-042). This isoindole-benzohydrazide derivative (**Figure 2E**) exhibited strong adjuvant activity when combined with 1 μg/mL COL, potentiating the antibiotic’s activity to terminate growth of the MCR-M6-expressing strain at all evaluated concentrations (**Figure 2D**). We assessed JNAL-042 for potential cytotoxicity against a human embryonic kidney cell line using a propidium iodide (PI) fluorescence membrane permeability assay.^36^ Results matched the solvent control, suggesting no notable impact on this cell line’s viability (**Figure 2E**) and warranting further biological characterization of this adjuvant.

**Figure 2.**
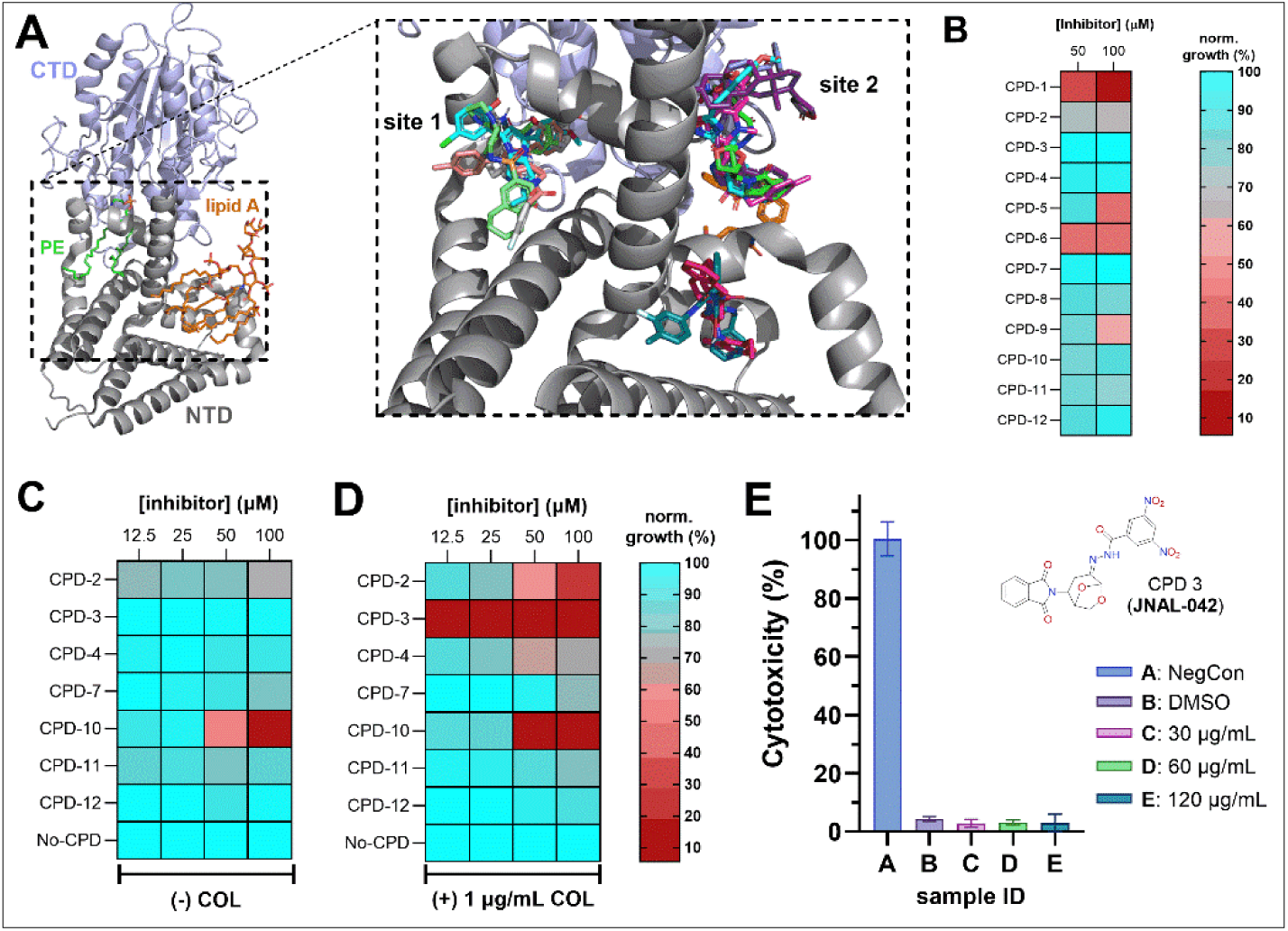
Combined computational and growth susceptibility screens identify candidate adjuvant for COL activity against MCR-mediated resistance. **A**) Cartoon representation of MCR-1 bound to its substrates, PE and lipid A, highlighting in silico virtual screening results for the top 12 prioritized compounds, which are predicted to bind to the PE (site 1) or lipid A (site 2) binding sites. **B**) Results from a dual-dose bacterial susceptibility assay evaluating test compounds for intrinsic growth-inhibitory activity in the absence of an antibiotic using the vector-carrying control strain, ASW-022. **C, D**) Results from four-dose growth susceptibility screens using an MCR-M6-expressing strain (ASW-018) in the absence (C) or presence (D) of 1 µg/mL COL. **E**) Dose-dependent cytotoxicity profiles for JNAL-042 against human embryonic kidney cells. The data represent 3 biological replicates (±SD).

### JNAL-042 is a promising potentiator of COL activity against MCR-mediated resistance

After identifying a candidate for our study, we conducted an extensive cheminformatics search for any existing biological activity footprint or projected protein targets. Still, we found no such reports in the ZINC20 database, PubChem, or using the Similarity Ensemble Approach.^38,40^ This search also reviewed available information based on structural similarities with known ligands, unsuccessfully.^38,40^ To characterize the adjuvant properties of JNAL-042, we first conducted checkerboard assays combining this compound with COL using the efflux-deficient strain (ASW-018) expressing the MCR-M6 variant. As observed in the initial 4-dose experiments (**Figure 2C, D**), JNAL-042 showed no growth-inhibitory activity when tested alone, with serial dilutions beginning at 48 µg/mL (100 µM) (**Figure S2**). Conversely, we recorded an MIC of 1-2 µg/mL for COL alone. In combination, 6 µg/mL JNAL-042 was sufficient to potentiate COL activity, resulting in a fractional MIC of 0.25 µg/mL, a ≥ 4-fold decrease that mimicked a sensitive phenotype (**Figure S2**). We complemented these experiments by determining the MIC profiles of COL and PMB with and without JNAL-042 against the VC, WT-MCR-1, and MCR-M6 in WT and TolC mutant *E. coli* strains. The obtained results corroborated trends observed in the checkerboard assays. For both antibiotics, WT MCR-1 caused a >2-fold MIC increase, with a smaller change in the efflux-deficient (RON-001) strain relative to the WT strain (RON-011) (**Table 1**). Conversely, the MCR-M6 variant caused a more drastic impact in both WT (ASW-020) and efflux-deficient (ASW-022) *E. coli*, inducing a 7 – 21-fold loss in activity for COL and PMB under our experimental conditions (**Table 1**). Importantly, adding JNAL-042 (12 µg/mL) to strains lacking the efflux pump was sufficient to overcome resistance conferred by both MCR-1 variants, fully restoring antibiotic activity (**Table 1**). In contrast, this dose of the adjuvant only partially rescued antibiotic activity in WT cells, lowering the MICs of COL and PMB by about 2-fold or 4.8-fold, respectively. This result suggested that JNAL-042 could be a substrate for the TolC efflux pump.

### JNAL-042 abolishes MCR-mediated resistance in time-kill assays

To assess the adjuvant’s ability to potentiate COL and PMB activity in actively growing cultures, we conducted time-kill kinetic growth experiments with the efflux-deficient strain ASW-021 (empty vector) and ASW-018 (p*mcr-M6*). For these experiments, we monitored cell growth using optical density (OD_600nm_) and cell death via colony-forming unit counts and the PI permeability assay. PI is a positively charged fluorescent dye that is permeable only to membrane-compromised bacterial cells and binds nucleic acids, enhancing its fluorescence to facilitate fluorescence-based cell-death quantification.^36,41^ Because polymyxin antibiotics cause membrane damage leading to lysis,^42,43^ we hypothesized that this dye would readily access cellular nucleic acids, providing an accurate method for quantifying cell death. For these assays, we typically used overnight cultures to inoculate fresh LB broth containing 5 µM PI, 0.5% w/v arabinose (expression-inducer), and kanamycin (50 µg/mL) for selection at a starting OD_600nm_ ∼0.05. We evaluated COL or PMB at concentrations ranging from 0–5 µg/mL, titrated across 96-well plates, followed by addition of strain ASW-021, ASW-018, or the latter supplemented with 12 µg/mL JNAL-042 in the bulk culture growth medium. We then incubated the plates and recorded growth (OD_600nm_) and PI fluorescence over 7 hours. We also conducted agar dilution plating assays to quantify viable cells at T = 0, 4, and 7 h post-assay initiation. The growth profiles of strain ASW-018 showed a 2- to 4-fold increase in the effective inhibitory concentration

(EIC) needed to completely arrest cell growth for PMB (0.63 μg/mL) and COL (2.5 μg/mL), respectively, relative to the control strain (0.63 µg/mL) (**Figure 3A, B; Figures S3 & S4**). Remarkably, adding 12 µg/mL JNAL-042 fully restored the activity of both antibiotics, resulting in an EIC equal to or lower than that needed to terminate growth of the sensitive strain, ASW-021 (**Figure 3C; Figures S3 & S4**). We also observed similar adjuvant effects of JNAL-042 when monitoring cell death by PI fluorescence. In this case, 0.63 µg/mL COL or PMB was sufficient to sterilize strain ASW-021 (**Figure 3D; Figures S3 & S4**). Strain ASW-018 showed a 2- to 4-fold increase in effective bactericidal concentration (EBC), and adding 12 µg/mL JNAL-042 to the culture reverted the EBC to control-strain levels for both antibiotics (**Figure 3D-F; Figure S3 & S4**). Further, we performed plating assays to determine cell viability counts with COL only (not PMB), as it elicited the largest observed increase (4-fold) in the dose required to inhibit growth. The results obtained corroborated the PI cell-death trends observed in the broth-microdilution experiments. In this case, 0.31 µg/mL COL sterilized strain ASW-021 but not ASW-018 after 4 h, with the latter showing resistance even at 1.3 µg/mL after 7 hours (**Figure 3G, H**). Likewise, adding 12 µg/mL JNAL-042 enabled 1.3 µg/mL COL to sterilize strain ASW-018, while achieving a significant (****p<0.0001) reduction in viable colonies at 0.16 and 0.31 µg/mL COL after 7 h (**Figure 3H, I**). We also conducted experiments to determine the frequency of spontaneous mutants resistant to COL (1 µg/mL) or a combination of COL and JNAL-042 (12 µg/mL), and calculated 3.7 × 10^-3^ for the antibiotic and 6.9 x 10^-5^ when the adjuvant was present, demonstrating about 2 orders of magnitude reduction. Colony counts for control samples treated with only JNAL-042 were similar to those of the drug-free control, suggesting that the potential emergence of resistance to the adjuvant was rare. These data establish the strong adjuvant properties of JNAL-042 in potentiating polymyxin antibiotic activity against MCR-M6-mediated resistance.

**Figure 3.**
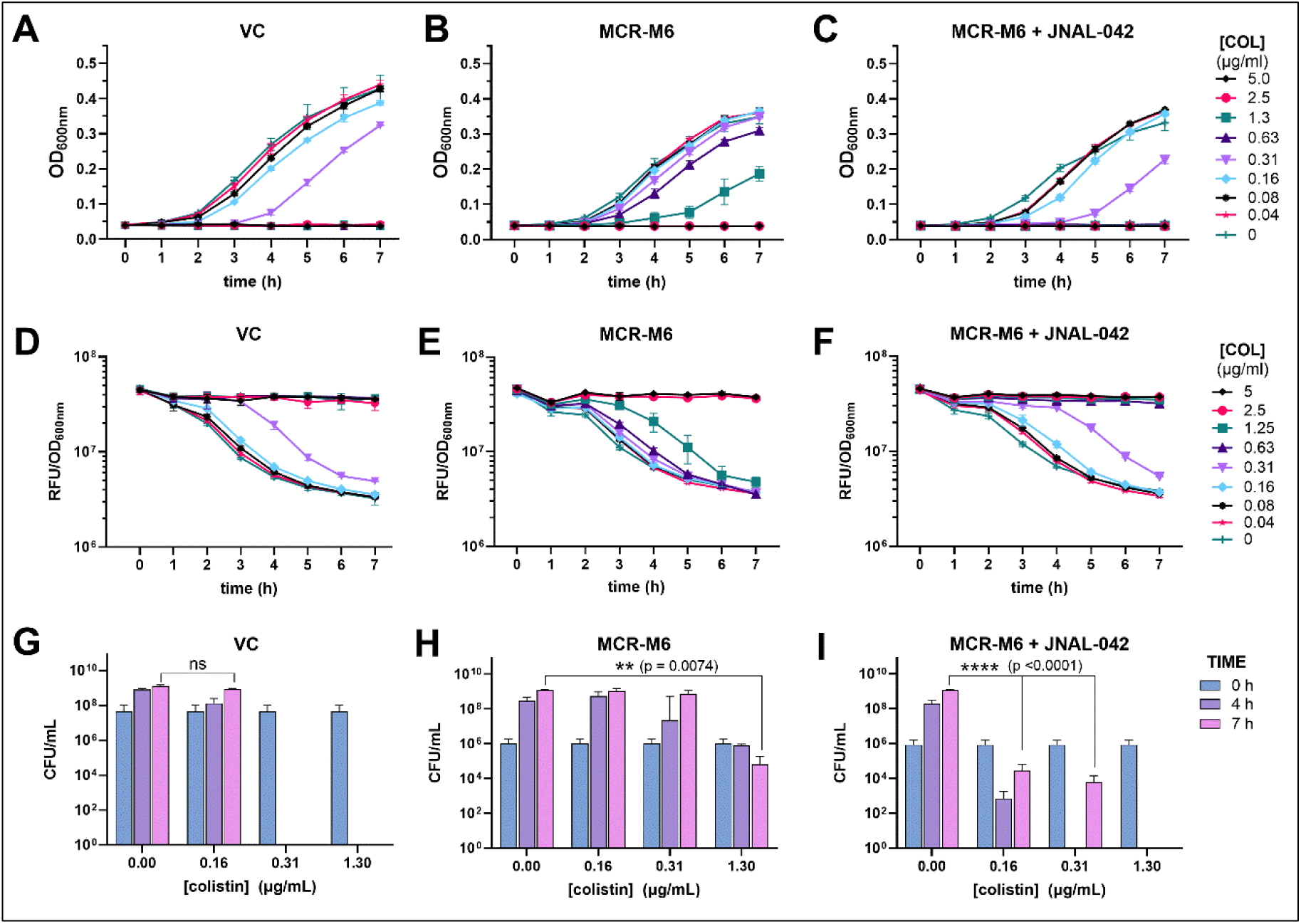
JNAL-042 potentiates COL activity to abolish MCR-1-mediated resistance in time-kill assays with efflux-deficient *E. coli* strains. The experiment used an engineered empty vector-carrying E. coli strain (VC; ASW-021; a COL-sensitive strain; **A, D,** & **G**) and one containing the gene encoding the MCR-M6 variant (a COL-resistant strain; **B-C, E-F,** & **H-I**). **A-C**) Time-dependent growth profiles of the VC and MCR-M6- expressing strains treated with varying COL concentrations monitored using cell density at OD_600nm_. **D-F**) Time-kill assay monitored using the PI assay, demonstrating the restoration of COL activity in the presence of JNAL-042 (12 µg/mL) to an effective bactericidal concentration (EBC) similar to that observed in the VC strain (0.63 µg/mL). **G-I**) Cell viability counts from solid agar dilution experiments assessing the bactericidal properties of COL. For graphs A−F, the data presented are averages from three technical replicates (±SD). Additional COL & PMB biological replicates are presented in **Figures S3 and S4**. For G-H, the data are presented as the mean from 3 biological replicates (+SD). Abbreviations: RFU – relative fluorescence units; CFU – colony-forming units.

### COL-JNAL-042 combination arrests cell growth in established resistance delayed-treatment experiments

Treatment of clinical cases involving bacteria-borne diseases typically occurs post-infection. Therefore, we designed experiments to mimic this scenario by monitoring time-dependent bacterial growth and cell death to assess whether JNAL-042 could potentiate COL activity to overcome established MCR-mediated resistance when added after enzyme production in bacterial cells. For these experiments, we used overnight cultures of strains ASW-018 and ASW-021 to inoculate bulk working stocks of LB broth supplemented with 0.5% arabinose and 5 µM PI to an OD_600nm_ of ∼0.1. We set up the cultures in 96-well plates and incubated them with shaking (325 rpm) at 37 °C. We monitored bacterial growth and PI fluorescence hourly and added COL at different concentrations, with or without JNAL-042 (24 µg/mL), 3 hours after assay initiation. We recorded growth and PI fluorescence for an additional four hours before quantifying the data. These experiments showed complete growth arrest of strain ASW-021 at an EIC of 2 µg/mL for COL (**Figure 4A; Figure S5**). A 2-fold increase in this EIC was required to inhibit growth when MCR-M6 was expressed, with no effect at 2 µg/mL (**Figure 4B; Figure S5**). Adding 24 µg/mL JNAL-042 abolished MCR-M6-mediated resistance, allowing 2 µg/mL COL to sustain growth arrest in the cultures (**Figure 4C; Figure S5**). We observed similar trends and corresponding EICs when assessing cell death using the PI fluorescence assay. Cells expressing the enzyme showed a 2-fold increase in the EBC, which reverted to control-vector levels in the presence of JNAL-042 (**Figure 4D-F; Figure S5**). These results highlight JNAL-042’s effectiveness in restoring COL activity in cells with established MCR-mediated resistance.

**Figure 4.**
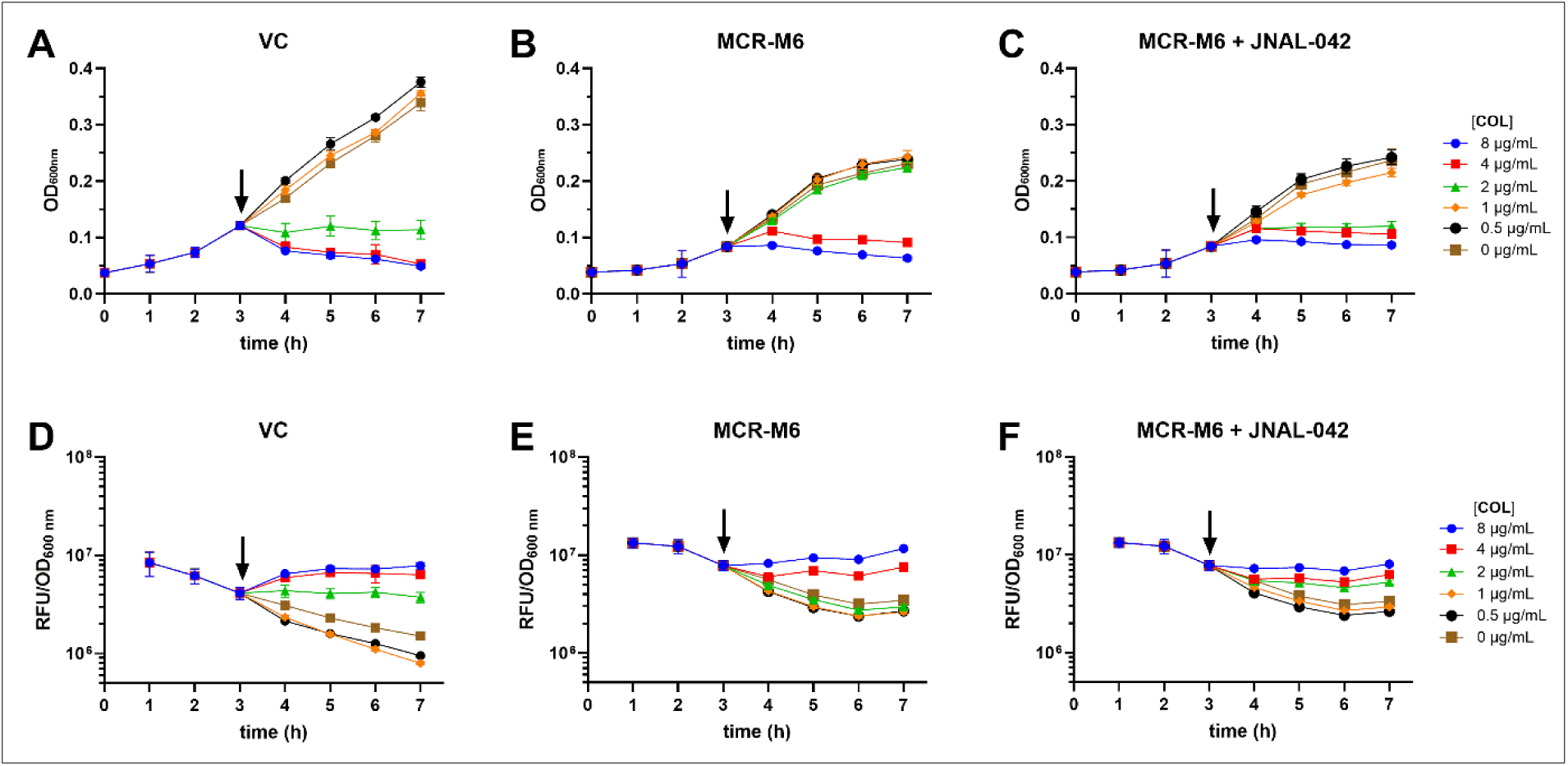
JNAL-042 potentiates COL activity against MCR-M6-mediated resistance in post-induction treatment time-dependent susceptibility experiments. **A-C**) Growth profiles monitored using cell density (OD_600nm_) for control (ASW-021) and MCR-M6-expressing strain (ASW-018), evaluating the effect of adding COL (A, B) or a combination of COL + 24 µg/mL JNAL-042 (C) 3 h (arrow) after inducing enzyme production. **D-F**) Time-dependent bacterial cell death profiles monitored over 7 h using the PI incorporation assays for the corresponding cultures in A-C. The data are averages from three technical replicates (± SD). See **Figure S5** for additional biological replicates. Abbreviations: VC – vector control; RFU – relative fluorescence units.

### COL-JNAL-042 combinations, but not PMB, require an efflux pump inhibitor to fully overcome MCR-M6-mediated resistance in WT *E. coli* cells

During our initial screens to identify candidate inhibitors of MCR variants in the *E. coli* model, we hypothesized that TolC efflux pump activity might prevent us from identifying effective candidates. Therefore, we used TolC mutant strains to screen and initially characterize our candidate adjuvants. To more closely mimic clinical infections, we conducted complementary experiments to assess the activity of COL-JNAL-042 combinations against WT *E. coli*. As demonstrated earlier in our MIC determination experiments, the presence of 12-24 µg/mL JNAL-042, a concentration range that acted synergistically with COL and PMB to fully restore their activity in the efflux-deficient strain, only partially restored their MICs in a WT background (**Table 1**). As hypothesized, this partial restoration could result from active cellular removal of JNAL-042 by the TolC efflux pump. Therefore, we argued that an efflux pump inhibitor could help to fully restore the COL and PMB activity when combined with the adjuvant. To test this hypothesis, we evaluated the activity of 1-(1-naphthylmethyl)piperazine (NMP), an efficient inhibitor of TolC with reported MIC values ranging from 400–800 µg/mL.^44,45^ We first determined 17 µg/mL as an effective NMP dose for our experiments. This concentration did not show growth-inhibitory activity even in the presence of colistin (**Figure S6**), in agreement with published reports.^44^

We conducted time-kill kinetic growth experiments using strains ASW-020 and ASW-022 to assess the effect of the antibiotic-JNAL-042 combination treatment on actively growing WT *E. coli* cells. These assays evaluated the activity of COL alone, COL+JNAL-042, and this combination in the presence of NMP at the concentrations highlighted above, while monitoring growth and cell death via PI fluorescence over 7 hours. We also performed parallel experiments to assess combinations of PMB-JNAL-042 with NMP. From these assays, COL had an EIC of 0.63 µg/mL in the control strain ASW-022 (**Figure 5A**), which increased to 5.0 µg/mL (8-fold) when expressing MCR-M6 (**Figure 5B**). Unlike the effect observed in similar assays involving efflux-deficient strains, adding 24 µg/mL JNAL-042 only partially restored COL activity, resulting in a 4-fold decrease in the EIC (1.25 µg/mL) required to arrest growth (**Figure 5C**). Interestingly, the addition of 17 µg/mL NMP to a COL-JNAL-042 combination treatment fully restored COL activity, yielding an EIC of 0.63 µg/mL (**Figure 5D**). Assessment of the COL-induced cell death using PI fluorescence showed similar trends, with the same COL growth EICs needed to kill cells in the corresponding assay treatments (**Figure 5E-H**). These results corroborated trends observed in the MIC data, demonstrating the need for an efflux pump inhibitor to fully restore COL activity in WT *E. coli* cells. On the contrary, time-kill assays with PMB revealed a surprising outcome. As shown by the MIC values (**Table 1**), PMB-JNAL-042 combinations similarly resulted in partial restoration of the antibiotic’s activity in WT cells expressing MCR-M6. For these kinetic assays, PMB displayed an EIC of 0.63 µg/mL in the control strain (**Figure 5I**). MCR-M6-induced resistance increased this value 4-fold (2.5 µg/mL) (**Figure 5J**). However, unlike the case with COL, combining PMB with JNAL-042 (24 µg/mL) fully restored the antibiotic’s activity, resulting in an EIC of 0.63 µg/mL (**Figure 5K**). Also, it is noteworthy that a cocktail including PMB, JNAL-042, and NMP further lowered the antibiotic’s EIC to 0.31 µg/mL (**Figure 5L**). This observation supports our argument that JNAL-042 is a substrate for TolC, and this efflux pump has a mild but negative impact on the COL-adjuvant combinatorial treatment. The data also suggest that higher doses of JNAL-042 might be sufficient to overcome the negative impact of efflux pumps.

**Figure 5.**
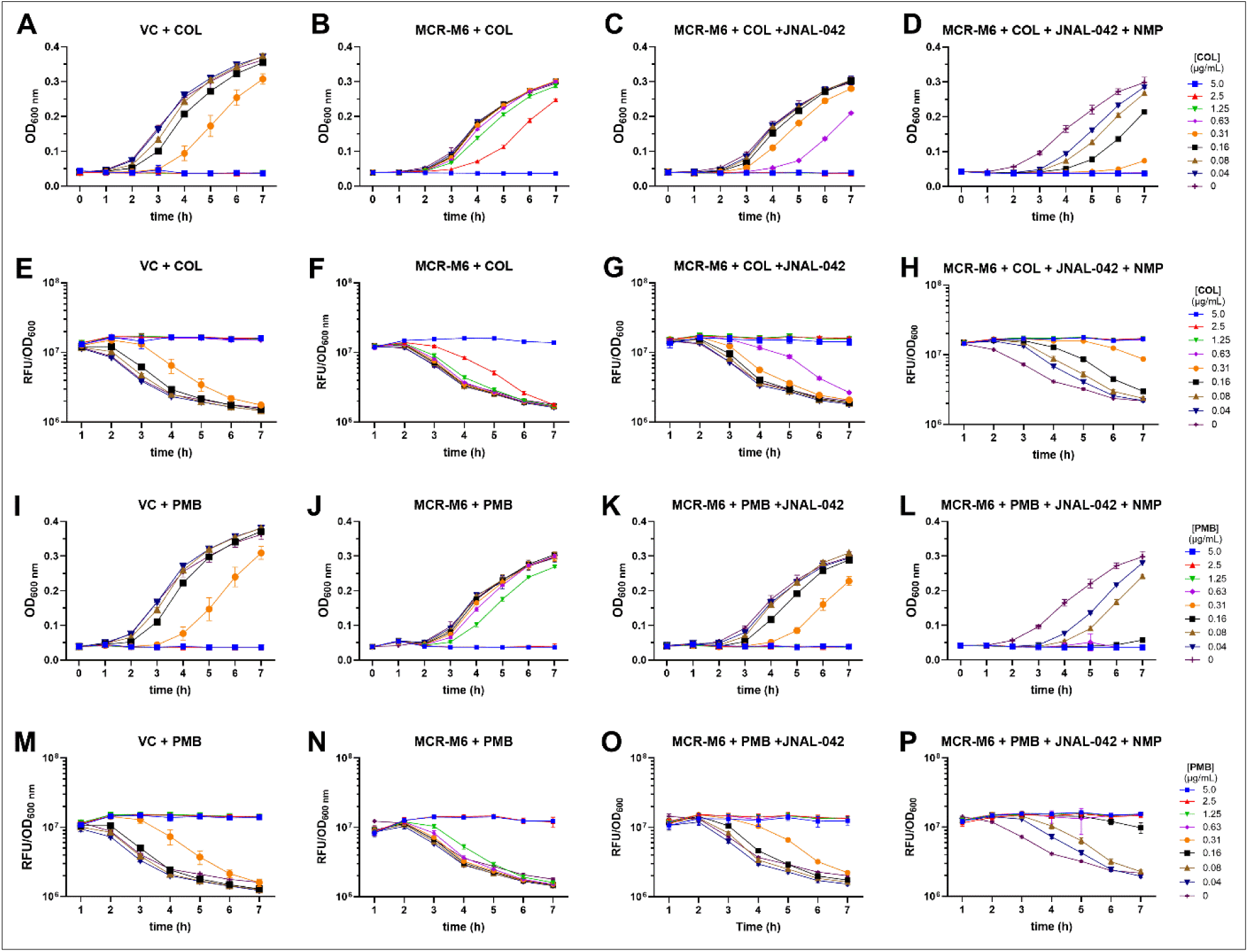
JNAL-042 suppresses MCR-M6-mediated resistance in WT *E. coli* cells. Results from experiments assessing the activity of COL in actively growing cultures by monitoring the time-dependent growth (OD_600nm_) (A-D) and cell death using a fluorescence-based PI incorporation reporter assay (E-H) in WT *E. coli* strains carrying an empty vector (ASW-022) or an MCR-M6-expression plasmid (ASW-020). **A, B, E, F**) Results obtained from monitoring COL activity. **C, G**) Activity profiles of COL-JNAL-042 (24 µg/ml) combination treatments in the presence of the enzyme. **D, H**) Data highlighting the activity of a treatment cocktail, including COL + JNAL-042 (24 µg/ml) + NMP (17 µg/ml), demonstrating complete restoration of COL activity. **I, J, M, N**) Results obtained from monitoring PMB activity. **K, O**) Activity profiles of PMB-JNAL-042 (24 µg/ml) combination treatments in the presence of the MCR-M6. **L, P**) Results demonstrating the activity of a treatment cocktail, including COL + JNAL-042 (24 µg/ml) + NMP (17 µg/ml), demonstrating complete restoration of COL activity. The data are averages of three technical replicates (±SD) from three biological replicates. See **Figures S7 & S8** for additional biological replicates. Abbreviations: VC: vector control; RFU: relative fluorescence units; NMP: 1-(1-naphthylmethyl)piperazine.

### JNAL-042 may block MCR activity by interfering with the second catalytic step involving PEtN transfer onto LPS

While ongoing structure-activity relationship and biochemical studies will help to uncover the mechanistic details of JNAL-042’s adjuvant activity, we used in silico docking studies to gain some insight into how this adjuvant may potentiate the activity of polymyxin antibiotics. In the initial virtual screens, we identified JNAL-042 by prioritizing compounds predicted to bind near the active- or substrate-binding sites. Subsequently, docking of this adjuvant localized it adjacent to the lipid A binding site, suggesting a potential mechanism involving inhibition of the second catalytic step (**Figure 6A-C**). To further investigate this hypothesis, we performed in vitro competitive binding experiments between JNAL-042 and *E. coli* LPS extracts with purified MCR-M6 using microscale thermophoresis.^46^ JNAL-042 had an estimated dissociation constant (K_d_) of 11.3 ± 1.9 µM (**Figure 6D**). However, preincubation of the enzyme with LPS, followed by addition of JNAL-042, abolished binding (K_d_ >250 µM), suggesting a competitive relationship between these species (**Figure 6D**). To investigate whether JNAL-042 might possess broad-spectrum activity against other MCR variants, we obtained their structures from the AlphaFold database or used this platform to predict them. Alignment of these variants demonstrated highly conserved structures (**Figure S9**). We then overlaid the docked structure of JNAL-042 with those of the variants and noted a high conservation of the adjuvant’s predicted binding pocket adjacent to that of lipid A (**Figure S9**). This observation suggests that the activity of JNAL-042 may be preserved across MCR variants. However, follow-up studies are needed to establish this projection. Therefore, based on the comprehensive computational and biological characterization data presented in this study, we propose a general model for this adjuvant’s mode of action in potentiating polymyxin antibiotic activity. In this model, JNAL-042 binds the enzyme in a location and conformation that may hinder the subsequent transfer of PEtN onto lipid A. This action prevents LPS modification, thereby promoting binding and membrane lysis by polymyxin antibiotics (**Figure 6E, F**). Taken together, our data support JNAL-042 as a promising adjuvant for combinatorial use with polymyxin antibiotics against MCR-mediated resistance.

**Figure 6.**
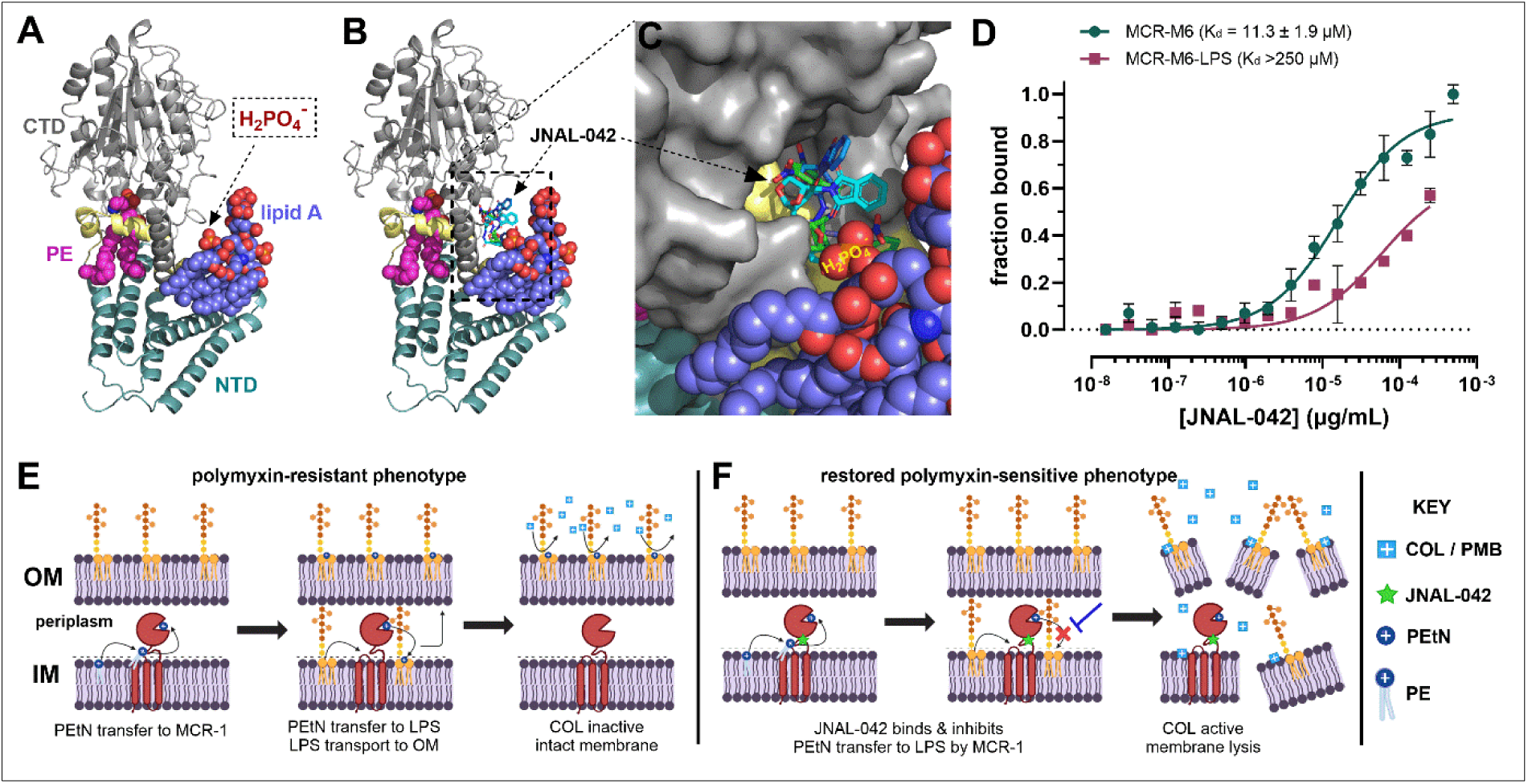
JNAL-042 may be interfering with PEtN transfer onto lipid A in the second catalytic step. **A**) Cartoon structure of MCR-1 bound to its substrates PE and lipid A (PDB ID: 9NWW) highlighting the phosphate functional group to be covalently modified by adding PEtN. **B**) Overlaid structures of MCR-1 in ‘A’ with docked structures of JNAL-042 depicting its predicted binding conformations in a pocket adjacent to lipid A. **C**) Close-up of the predicted binding pocket of JNAL-042 highlighting its proximity to a lipid A phosphate group, typically targeted for modification by the enzyme in the second catalytic step. **D**) In vitro binding profiles between MCR-M6 and JNAL-042 (circles) or this adjuvant and a premixed sample of the enzyme with 0.16 mg/ml *E. coli* LPS extracts (squares). Data represent the mean (±SD; n = 3). **E**) Cartoon schematic of the mechanism of MCR-1-mediated resistance to polymyxin antibiotics, highlighting the covalent modification of LPS preventing COL from binding to cause membrane lysis. **F**) Proposed mode of action of JNAL-042, which is hypothesized to bind adjacent to lipid A, preventing the covalent modification of LPS, promoting COL/PMB activity in the polymyxin-resistant strain.

## DISCUSSION

Persistent multidrug-resistant bacteria are forcing the prioritization of last-resort antibiotics, including polymyxins, to treat life-threatening clinical infections. The rapid spread and evolution of the *mcr* gene raise serious concerns, as it threatens this life-saving option for combating deadly pathogens, including carbapenem-resistant Enterococcus species, with polymyxin antibiotics. Some studies have made valid attempts to target MCR-mediated resistance using CADD.^27–30,47^ However, most prioritize computational approaches or lack accompanying comprehensive biological characterization data for identified potentiators, calling for renewed efforts to address this deficiency. In this study, we identified and characterized the biological activity of a promising potentiator of polymyxin antibiotic activity against MCR-1-mediated resistance in *E. coli*. Following phenotypic validation of our model strains, we employed computational virtual screens to demonstrate the druggability of MCR-1 with compounds from a natural product library. For this study, we prioritized prospective inhibitors projected to bind within or adjacent to the substrate-binding sites of the enzyme (**Figure 2**), envisioning the identification of competitive inhibitors. This strategy led to the 12 candidate compounds used in our initial screens. First, we identified and eliminated compounds with intrinsic antibiotic activity, as they may target essential bacterial components or pathways, which was beyond the scope of our study. These screens successfully identified JNAL-042 as our primary adjuvant, whose biological characterization using kinetic growth assays validated its ability to abolish MCR-1-mediated resistance to polymyxin antibiotics at low concentrations (12 µg/mL), fully reverting the enzyme-expressing strain to a sensitive phenotype. Since COL is a bactericidal antibiotic whose mode of action typically results in cell death via membrane lysis, we also investigated the adjuvant’s ability to restore this crucial property. Accordingly, we accompanied the growth-inhibition experiments with time-kill assays, monitored using fluorescence-based PI incorporation and cell viability counts. Our collective results corroborated the adjuvant’s ability to potentiate COL activity, revealing a significant (****p < 0.0001) reduction in viable cell count even at 0.16 µg/mL of the antibiotic – a concentration that was ineffective in a control strain lacking the *mcr-M6* gene (**Figure 3G-I**). From a different perspective, patients typically seek medical assistance after experiencing symptoms from an infection. Therefore, we also assessed JNAL-042’s ability to elicit a similar adjuvant effect on COL in experiments mimicking post-infection treatment of a resistant strain, with antibiotic or antibiotic-adjuvant combinations added 3 hours after inducing MCR-M6 expression. Notably, while these kinetic time-kill experiments required a higher dose of COL to arrest growth or kill cells, JNAL-042 achieved comparable results, reverting the MCR-carrying strain to a sensitive phenotype. This result highlights JNAL-042’s effectiveness in abolishing established MCR-1-mediated resistance in actively growing cells, thereby restoring the antibiotic’s activity (**Figure 4**).

A critical consideration in drug discovery efforts involving bacteria is the presence of efflux pumps, which may affect the efficacy of candidate compounds. Because our initial validation studies for JNAL-042 used efflux-deficient strains, we evaluated this adjuvant’s ability to suppress MCR-mediated resistance in WT *E. coli* cultures to better mimic clinical isolates. MCR-M6 expression increased the effective bacteriostatic and bactericidal concentrations of COL 8-fold in kinetic time-kill assays (**Figure 5**). JNAL-042 (24 µg/mL) partially restored COL activity, achieving a 4-fold reduction in the EIC after 7 hours. Accordingly, we hypothesized that this suppressed activity may have resulted from active efflux of the adjuvant. Indeed, a treatment cocktail containing COL, JNAL-042, and NMP, a known inhibitor of TolC activity,^44,45^ successfully restored the antibiotic’s activity to similar levels needed to inhibit growth or kill empty-vector-carrying cells (**Figure 5**). The phenomenon observed with PMB, which, when combined with JNAL-042 (24 µg/mL), did not require NMP for full restoration of its activity, was puzzling. However, we note that MCR-M6 resulted in only a 4-fold loss of activity for PMB, compared with 8-fold for COL. This lower change in the EIC/EBC of PMB could explain the observed phenomenon, as the presence of the adjuvant only caused a 4-fold decrease in these values – similar changes were observed for COL-JNAL-042 combinations. These data highlight the need for an efflux pump inhibitor when assessing COL-JNAL-042 combinations against WT clinical isolates. They also call for follow-up experiments to evaluate higher adjuvant doses, given that the adjuvant demonstrated no cytotoxic side effects against HEK 293 cells at 5x the dose used in these experiments.

Importantly, despite the absence of detailed biochemical evidence for the mechanism of JNAL-042, its lack of intrinsic antibiotic activity suggests that its role in potentiating polymyxin activity is solely tied to the MCR resistance-conferring mechanism. Indeed, the competitive binding experiments with *E. coli* LPS extracts to purified MCR-M6 demonstrated a >25-fold loss in affinity for the adjuvant (**Figure 6D**). This observation suggests that JNAL-042 competes for the LPS binding site, consistent with our proposed mode-of-action model. In this model, which is partially based on computational predictions, JNAL-042 binding would prevent PEtN transfer by MCR-M6 to the adjacent phosphate group of lipid A. While this hypothesis warrants further investigation, it supports an intriguing strategy to inhibit MCR-1 activity by blocking the second catalytic step. Moreover, structural alignment of MCR-1 – 10 variants with the docked structure of JNAL-042 (**Figure S9**) suggests that this adjuvant could possess broad-spectrum properties against strains displaying this form of resistance. Nonetheless, the computational and biological data presented in this study provide compelling evidence demonstrating that JNAL-042 is a promising adjuvant for potential use with polymyxin antibiotics against MCR-mediated resistance - a discovery with significant clinical implications.

### Limitations of the study

This study characterized the biological activity of an isoindole-benzohydrazide derivative, JNAL-042, which potentiates polymyxin antibiotic activity against MCR-1-mediated resistance. While it includes sufficient biological data to support the key findings, limitations include a lack of detailed biochemical data to validate the proposed mechanism of action of the adjuvant. Likewise, the data presented originated from laboratory strains of *E. coli*. Validating the potential clinical application of this adjuvant would benefit from evaluating clinical isolates exhibiting MCR-mediated resistance. Despite these limitations, our findings provide perhaps the first comprehensive characterization of an adjuvant with validated activity to restore polymyxin antibiotic activity against MCR-1-mediated resistance. Furthermore, ongoing structure-activity relationship and biochemical studies assessing JNAL-042 analogs should provide the full scope of its mechanism of action, with in vivo studies on the horizon.

## Resource availability

### Lead contact

Further information and requests for resources and reagents should be directed to and will be fulfilled by the lead contact, John N. Alumasa.

### Materials availability

Further information and requests for resources and reagents should be directed to and will be fulfilled by the lead contact, John N. Alumasa.

### Data and code availability

- Any additional information required to reanalyze the data reported in this paper is available from the lead contact upon request.
- This paper does not report original code.

## Acknowledgements

This work was supported by faculty research startup funds to J.N.A. from the Department of Chemistry and Biochemistry at Miami University, Oxford, Ohio.

Schematics were prepared using BioRender.com (https://BioRender.com/l38ojdl; https://BioRender.com/kqr8meo).

## Author contributions

Conceptualization, J.N.A.; A.S.W. performed all experiments involving the MCR-M6 variant. R.O.N. performed all experiments involving the WT MCR-1. A.S.W. drafted the methods section. J.N.A. drafted and wrote the manuscript. A.S.W., R.O.N., and J.N.A. reviewed the final manuscript.

## Materials & Methods

### Strain & Plasmid Table

| Strain | Description | Source |
| --- | --- | --- |
| BW25113 (WT) | *E. coli* strain with *araBAD* deletion to inhibit arabinose metabolism | ATCC<sup>#</sup> |
| BW25113Δ*tolC::cat* | BW25113 (WT) with *tolC* (efflux pump) deletion, replaced by the chloramphenicol acetyltransferase resistance gene. | Ref (35) |
| BL21(DE3) | *E. coli* protein expression strain. | NEB<sup>\$</sup> |
| RON-001 | BW25113Δ*tolC::cat* strain containing plasmid pRON-001 | This study |
| RON-011 | BW25113 (WT) strain containing plasmid pRON-001 | This study |
| ASW-018 | BW25113Δ*tolC::cat* strain containing plasmid pASW-001 | This study |
| ASW-020 | BW25113 (WT) strain containing plasmid pASW-001 | This study |
| ASW-021 | BW25113Δ*tolC::cat* strain containing a pMyBADC-kan vector. | This study |
| ASW-022 | BW25113 (WT) strain containing a pMyBADC-kan vector. | This study |
| ASW-023 | BL21(DE3) strain containing plasmid pASW-002 | This study |
| **Plasmid** |  |  |
| pMyBADC-kan | Expression vector used for the controlled production of MCR-M6 (*P<sub>BAD</sub>* promoter-AraC-regulator; arabinose - induction). Contains a kanamycin resistance marker. | Addgene |
| pASW-001 | pMyBADC-kan vector containing the *E. coli mcr1-m6* gene for regulated expression of MCR-M6 in cell-based assays. | This study |
| pASW-002 | pET28a vector containing the *E. coli mcr1-m6* gene for the expression and purification of the MCR-M6 variant. | GenScript |
| pRON-001 | pMyBADC-kan vector containing the *E. coli mcr-1* gene for regulated expression of WT MCR-1 in cell-based assays | This study |

<sup>#</sup>ATCC – American Type Culture Collection; <sup>\$</sup>NEB – New England Biolabs

### Construction of MCR-1 and MCR-M6 resistance model strains

The *E. coli* WT *mcr-1* gene sequence (UniProt: A0A0R6L508) was manually edited to create the *mcr-M6* variant strain containing amino acid substitutions P188A and P195S.^31^ Sequenced plasmid constructs containing these genes were obtained from GenScript (New Jersey). WT *mcr-1* was obtained as a plasmid construct with the gene cloned into the pET21a(+) vector using the 5’-NheI and 3’-HindIII sites. The gene was PCR-amplified using primers containing 5’-EcoR1 and 3’-SalI restriction sites (forward primer: 5’-ATATATGAATTCATGATGCAGCATACCAGC, reverse primer: 5’-TATATAGTCGACTTAGCGAATAAAGGCGGT), before being subcloned into the pMyBADC-kan vector to generate plasmid pRON-001. The *mcr-M6*-carrying plasmid was obtained in a pET28a vector and directly subcloned into the pMyBADC-kan expression vector using the 5’-NheI/3’-SalI restriction sites, resulting in plasmid pASW-001. The final plasmid constructs were confirmed by Sanger sequencing. These plasmids were initially transformed into DH5α cells and finally into BW25113Δ*tolC::cat* or WT BW25113 cells, yielding strains RON-001, RON-011, ASW-018, and ASW-020 (see strain table for details). The pMyBADC-kan empty vector was also transformed into BW25113Δ*tolC::cat* or WT BW25113 cells to generate control strains, ASW-021 and ASW-022, lacking the resistance-conferring enzymes.

### Determination of minimum inhibitory concentrations (MICs)

Overnight cultures (5 mL) of corresponding strains were diluted 1:500 in lysogeny broth (LB) containing 50 μg/mL kanamycin and 0.2% arabinose and grown to OD ∼1.0. Bulk working cultures were then prepared by diluting the cultures in fresh LB containing 50 μg/mL kanamycin and 1% arabinose (final 2X OD_600nm_ 0.004). Serial dilutions of test compounds or antibiotics at 2X the intended final concentration were prepared in 50 µL LB in a 96-well plate, and 50 µL of the bulk working culture was added to each well for a final starting OD_600nm_ of 0.002. The assay plates were incubated at 37°C for 16–18 h before recording MIC values, which were determined as the lowest concentration with no visible growth following a visual inspection. Absorbance readings of the microplates were recorded using a SpectraMax iD5 plate reader (Molecular Devices) and normalized to “drug-free” controls to generate dose-response plots using GraphPad Prism. Three biological replicates were performed for each. For assays using fixed JNAL-042 concentrations, this adjuvant was added to the bulk working cultures at 2x the desired final concentration.

### In Silico virtual screening and molecular docking of JNAL-042

Computational virtual screens were conducted using the AutoDock Vina software. The cryo-EM structure of MCR-1 (PDB ID: 9NWW) bound to its two substrates (PE and Lipid A) was used for these screens. Note: The bound substrate structures were removed during the docking event. Structure-Data files of 10,191 compounds were retrieved as a compressed file from a virtual natural product library from the ZINC20 database and loaded into the docking software. An Open Babel platform integrated into the docking software was used to perform energy minimizations of the structures, followed by file conversion into the required pdbqt format. Docking screens were performed with a search grid covering the entire MCR-1 structure, and the obtained binding energies were used to select initial candidates, focusing on compounds predicted to have binding energies (cut-off: -8.0 kcal/mol). This screening identified 1,682 compounds of interest.

Subsequently, the SwissADME (absorption, distribution, metabolism, and excretion) online structural evaluation software was used for secondary screening to identify candidates with suitable predicted pharmacokinetic and drug-like properties. The top 12 compounds based on this ranking were selected and purchased for our proof of concept study. Docking of JNAL-042 was conducted in a similar manner, and several predicted conformations were used for analysis.

### Preliminary screens to identify adjuvants against MCR-M6-mediated resistance

The dual-dose and quad-dose cell-based screens involving the computationally identified mini-library of prospective adjuvants were performed using similar setup protocols as those described for the MIC assays above. When assessing the compounds alone, a 2X bacterial culture (OD_600nm_ 0.004) of a corresponding strain supplemented with 50 μg/mL kanamycin and 1% arabinose (where necessary) was added to serially diluted test compounds in 96-well microplates. When required, 2 µg/mL of colistin was added to the bulk culture (final concentration 1 µg/mL). The plates were incubated at 37°C for 16 – 18 h, and the growth data were analyzed by obtaining the optical densities (OD_600nm_), which were normalized to the drug-free control samples. Heat maps of the resulting data were generated using GraphPad Prism.

### Combinatorial checkerboard assays to assess COL-JNAL-042 synergy

A 5 mL overnight culture of strain ASW-018 grown in LB supplemented with 50 µg/mL kanamycin was diluted to a 2X working stock in the same medium containing 1% arabinose to an OD_600nm_ 0.004. Colistin or JNAL-042 dilutions (2-fold with 50 µL volume transfer) were performed in a 96-well plate using LB containing 50 µg/mL. Colistin was serially diluted across the plate (columns 1-9, rows B-H), starting at 4X the desired final concentration, and JNAL-042 down the plate (columns 2-9, rows A-H), beginning at 2X the desired final concentration. Columns 10 and 11 were used to determine MICs of Colistin and JNAL-042, respectively. Drug-free DMSO control samples were included in column 12. The 2X diluted culture (50 µL) was added to each well (final OD_600nm_ 0.002), and the plate was incubated at 37°C for 18 h. MIC or fractional MIC values were determined by visual inspection of the plate to identify the lowest concentration without visible growth. OD_600nm_ values of the samples were also obtained using a SpectraMax® iD5 plate reader (Molecular Devices), and heat maps were generated with GraphPad Prism. Three biological replicates were conducted, and the data were averaged to obtain the mean values. The highest final concentrations of COL and JNAL-042 used were 4 μg/mL and 48 μg/mL, respectively.

### Evaluation of cytotoxicity against Human Embryonic Kidney Cells

Human Embryonic Kidney Cells (HEK-293; ATCC CRL-1573; batch number 70071258) were cultured in Dulbecco’s Modified Eagle Medium (Gibco) supplemented with 10% fetal bovine serum and 1X GlutaMAX (Gibco). Sterile black-clear bottom 96-well tissue culture treated plates (Corning) were seeded with 1×10^5^ cells per well and placed in a 5% CO_2_ incubator at 37°C to allow cell attachment. JNAL-042 dilutions or DMSO controls were prepared in the same medium, which was pre-warmed to 37°C. These samples were used to replace media in the plates with JNAL-042 concentrations ranging from 30 µg/mL to 120 µg/mL. 2.5% DMSO was used as a control, and several drug-free wells were set aside as lysis control samples. After 24-hour incubation, 20 µL lysis reagent (10 mM Tris-Cl (pH 7.4), 150 mM NaCl, 1 mM EDTA, 1% Triton X-100) was added to designated lysis-control wells, and the plate was incubated for 10 minutes at 37°C, 5% CO_2_. Propidium iodide diluted in DMEM was then added to all the wells (5 µM final concentration), and the plate was incubated for an additional 20 minutes before measuring the fluorescence signal (λ_ex_ 535 nM, λ_em_ 615 nM) on a SpectraMax iD5 (Molecular Devices). The fluorescence data were normalized to the lysis control (representing 100% lysed cells) to quantify the relative toxicity. Three biological replicates were performed, and the mean values (± standard deviation) were used to generate the graph using GraphPad Prism 11.1.0.

### Time-kill assays and determination of colony-forming units

Note: incubation was performed at 37°C in a dry-air incubator shaking at 300 rpm. 50 µL of LB was added to each well of a 96-well plate. For strain ASW-018, an additional 46.8 µL of LB was added to column A, 3.2 µl sterile water was added to wells A1 and A12, and 3.2 µL of 10 mg/mL COL (final 2X concentration 320 µg/mL) was added to wells A2-10 to wells A2-10. For strain ASW-022, an additional 46 µL of LB was added to column A, 4 µl sterile water was added to wells A1 and A12, and 4 µL of 1 mg/mL COL (final 2X concentration 40 µg/mL) was added to wells A2-10. 2-fold serial dilutions were performed from row A through H, and 50 µL was discarded after the last well. A 2X bulk culture (OD_600nm_ = 0.05) of each strain was prepared with 50 µg/mL kanamycin, 10 µM PI fluorescent dye, and 1% arabinose, and 50 µL aliquots were added to each well for its respective plate. For plates including JNAL-042, JNAL-042 at 2X the final concentration was added to the bulk culture (2X concentration 24 µg/mL). Data at the T = 0 time point were obtained by recording the OD_600nm_ and PI fluorescence (λ_ex_ 535 nM, λ_em_ 615 nM) using the SpectraMax® iD5 plate reader (Molecular Devices). Each plate was then incubated for 7 h while recording the cell density and fluorescence readings every hour. The time-dependent growth curves and time-kill profiles were analyzed using GraphPad Prism. For the latter, the fluorescence values were normalized by dividing by the cell density of the corresponding sample well before quantification. Determination of the colony-forming units (CFU/mL) was performed with the same samples by extracting aliquots (5 µL) at 0, 4, and 7-hour time points. Depending on the cell density recorded in the culture wells, the extracted volumes were diluted as follows: cultures with an OD_600nm_ ≤ 0.15 were diluted 10^4^ fold, while those with a density > 0.15 were diluted 10^5^ fold using sterile water. Aliquots (50 µL) from these dilutions were then plated onto LB-Kan plates. The plates were incubated for 24 h, and colonies were counted visually, factoring in the dilution factors to determine the CFU/mL. Three biological replicates were conducted for these assays, and the data were analyzed using GraphPad Prism. Time-kill assays involving WT *E. coli* strains ASW-020 and ASW-022 were conducted in a similar manner, and JNAL-042 and NMP were assessed at 24 µg/mL and 17 µg/mL (final), respectively, in the corresponding samples.

### Delayed time-kill assays

Note: incubation was performed at 37°C in a dry-air incubator shaking at 300 rpm. LB (50 μL) containing 50 µg/mL kanamycin was added to each well of a 96-well plate. A 2X bulk culture (2X OD_600nm_ = 0.05) of ASW-018 was prepared with 50 µg/mL kanamycin, 10 µM PI fluorescent dye, and 1% arabinose, and 50 μL was transferred to each well of the plate. The T=0 time point was obtained by recording the OD_600nm_ and PI fluorescence (excitation at 535 nm and emission at 615 nm) using the SpectraMax® iD5 plate reader (Molecular Devices). The plate was then incubated for 3 h while monitoring the cell density and fluorescence readings every hour. After the T = 3 time point was recorded, serial dilutions of colistin were performed in either DMSO or DMSO containing JNAL-042 such that after the transfer of 5 μL of the solution to the culture-containing 96-well plate, each well would contain one of six concentrations of colistin ranging from 0.16 μg/mL to 5 μg/mL, and wells with the adjuvant would contain 24 μg/mL JNAL-042. Colistin dilutions were added to wells C1-C6 through H1-H6 (total of 12 technical replicates), and colistin dilutions containing JNAL-042 were added to wells C7-C12 through H7-H12 (total of 12 technical replicates). Plain DMSO was added to wells A1-A6 through B1-B6, and DMSO containing JNAL-042 was added to wells A7-A12 through B7-B12. The plate was then incubated for an additional 4 h while monitoring the cell density and fluorescence readings every hour. The time-dependent growth curves and time-kill profiles were analyzed using GraphPad Prism. For the latter, the fluorescence values were normalized by dividing by the cell density of the corresponding sample well before quantification. Three biological replicates were conducted for these assays.

### Combinatorial assessment of colistin, JNAL-042, and 1-(1-naphthylmethyl)-piperazine

Overnight cultures of strains ASW-020 and ASW-022 were diluted 1:500 in lysogeny broth containing 50 μg/mL kanamycin and 0.2% arabinose and grown to OD_600nm_ ∼1.0. Bulk working stocks (2X) were prepared by diluting these cultures in LB containing 50 μg/mL kanamycin, 24 μg/mL JNAL-042, and 1% arabinose (OD_600nm_ 0.004). LB broth containing 2X the intended concentration of colistin (final concentrations 0 μg/mL, 0.3 μg/mL, and 1.25 μg/mL) was prepared, and 50 μL was added to the respective lanes in a 96-well plate, along with an additional 46 μL in column A. 4 μL of the efflux pump inhibitor was added to column A, and a 2-fold serial dilution was performed across the plate. The 2X bulk working culture was added to each well for a final starting OD_600nm_ of 0.002. Absorbance readings of the plates were recorded using a SpectraMax iD5 plate reader (Molecular Devices) and normalized to drug-free controls to generate plots using GraphPad Prism. Three biological replicates of each experiment were obtained.

### Determination of the frequency of spontaneous mutations

A saturated culture of ASW-018 was diluted by transferring 30 μL of the overnight culture into 3 mL of fresh LB containing 50 μg/mL kanamycin and 0.2% arabinose and grown to OD_600nm_ ∼1.5. This culture was diluted to OD_600nm_ 1.0 in sterile water, and diluted 10^2^ and 10^4^ fold in sterile water. LB plates containing 50 μg/mL kanamycin, 1% arabinose, and colistin (1 μg/mL), JNAL-042 (12 μg/mL), or a combination of both compounds were prepared. Then, 50 μL aliquots of cells diluted 10^2^-fold were plated on pre-warmed LB agar plates containing both drugs, and 50 μL aliquots of cells diluted 10^4^-fold were added to control LB agar plates and plates containing colistin or JNAL-042. The plates were incubated at 37°C for 16 h. Colony counts were converted to CFU/mL values, and the frequency of resistance was calculated by dividing the CFU/mL values from each sample by that of the drug-free control plate. Three biological replicates were performed, and the data were averaged to obtain the final frequency of spontaneous mutations.

### Expression and purification of recombinant MCR-M6

Expression and purification of recombinant MCR-M6 was performed according to Sun, *et al*.,^48^ with some modification. Six overnight cultures of strain ASW-023 were used to prepare six 25 mL seed cultures in terrific broth (TB) and grown at 37°C with shaking at 250 rpm to OD_600nm_ ∼1.0. Each culture was used to inoculate 1 L of TB containing 50 μg/mL kanamycin and grown at 37 C, 150 rpm, to an OD_600nm_ ∼ 0.8. Induction was done overnight with 1 mM IPTG for 16 h at 18°C, shaking at 150 rpm. Cells were harvested by centrifugation at 4°C and 12,390 x g, frozen with liquid nitrogen, and stored at -80°C until further use. For purification, the cell pellet was thawed on ice and resuspended in lysis buffer (20 mM HEPES pH 7.4, 50 mM NaCl, 20 mM imidazole, 1 mM PMSF, 0.1 mg/mL lysozyme, 5 mM β-mercaptoethanol (BME). The cells were then lysed by sonication and clarified at 39,200 x g (4 °C). The lysate was filtered using a 0.45 μM filter and centrifuged for 1 hour at 80,000 x g (4 °C). The resulting pellet was solubilized overnight at 4 °C in PBS containing 2% DDM, and centrifuged at 30,624 x g for 30 minutes. The supernatant was passed through a 5 mL HisTrap HP Ni-NTA column (Cytiva) equilibrated with binding buffer (20 mM HEPES pH 7.4, 50 mM NaCl, 20 mM imidazole, 5 mM BME, 0.03% n-Dodecyl-β-D-maltoside (DDM) and washed with 5 CV of the same buffer. The enzyme was eluted from the column in 1 mL fractions using the AKTA Pure (Cytiva) with elution buffer (20 mM HEPES pH 7.4, 50 mM NaCl, 500 mM imidazole, 0.023% DDM, 5 mM BME), and each fraction was analyzed using SDS-PAGE. Fractions containing MCR-M6 were pooled and concentrated with a 10,000 MWCO filter to 2.5 mL. The protein was buffer-exchanged into storage buffer (20 mM HEPES pH 7.4, 50 mM NaCl, 0.03% DDM, 5 mM BME, 20% glycerol) using a PD10 desalting column (Cytiva).

### Binding affinity assessment using Microscale Thermophoresis (MST)

Purified 6xHis-N-terminal-tagged MCR-M6 (400 nM) was combined with 400 nM NTA-Atto 647 dye (Supelco) in PBS-T (phosphate-buffered saline; 137 mM NaCl, 2.7 mM KCl, 10 mM Na_2_HPO_4_, 1.8 mM KH_2_PO_4_, pH 7.4, containing 0.05% Tween-20). This sample was protected from light and incubated for 30 minutes at room temperature, followed by centrifugation at 15,000 x g for 10 minutes at 4°C. The supernatant was then transferred to a new microfuge tube. An aliquot of this labeled protein sample (5 μL) was added to 5 μL of serially diluted JNAL-042 samples (2X concentrations ranging from 0.015 μg/mL to 481.4 μg/mL). For competition experiments, 0.32 mg/mL LPS (final 0.16 mg/mL) was preincubated with the labeled protein on ice for 5 minutes, then this solution was added to JNAL-042 dilutions as described above. These mixtures were loaded onto Monolith capillaries (NanoTemper Technologies), and binding was assessed using microscale thermophoresis on a Monolith X (NanoTemper Technologies). The data were analyzed, and the dissociation constants of JNAL-042 were determined using GraphPad Prism. The K_d_ values from at least three independent experiments were averaged to provide the mean (± SD).

### Statistical data analysis

Statistical analysis was performed using R (version 4.6.1) with the tidyverse and emmeans packages.^49,50^ Data were analyzed using one-way analysis of variance (ANOVA) followed by post-hoc multiple comparisons using Dunnett’s test. A *P* value of 0.05 or less is considered statistically significant.

### Structural alignments of MCR variants

Sequences of MCR variants 2-10 (UniProt: A0A1C3NEV1, A0ABD7EMD5, A0A2I6AXF4, A0A2H4N789, A0A220T317, A0A2I7QFI7, A0A5K6TI91, A0A5B9AR78, A0A516EKB3) were obtained from UniProt. The predicted structures of each variant were obtained from the AlphaFold Protein Structure Database, and that of MCR-M6 was predicted using AlphaFold 3. Multiple structural alignment with MCR-1 (PDB: 9NWW) was performed and visualized using PyMOL.

## References

1. Chandrasekhar, D., Joseph, C.M., Parambil, J.C., Murali, S., Yahiya, M., and K, S. (2024). Superbugs: An invisible threat in post antibiotic era. Clin. Epidemiol. Glob. Health 28, 101499.

2. Halawa, E.M., Fadel, M., Al-Rabia, M.W., Behairy, A., Nouh, N.A., Abdo, M., Olga, R., Fericean, L., Atwa, A.M., El-Nablaway, M., et al. (2024). Antibiotic action and resistance: updated review of mechanisms, spread, influencing factors, and alternative approaches for combating resistance. Front. Pharmacol. 14, 1305294.

3. Naddaf, M. (2024). 40 million deaths by 2050: toll of drug-resistant infections to rise by 70%. Nature 633, 747–748.

4. Sun, S. (2025). Emerging antibiotic resistance by various novel proteins/enzymes. Eur. J. Clin. Microbiol. Infect. Dis. 44, 1551–1566.

5. Egorov, A.M., Ulyashova, M.M., and Rubtsova, M.Y. (2018). Bacterial enzymes and antibiotic resistance. Acta Naturae 10, 33–48.

6. Schaenzer, A.J., and Wright, G.D. (2020). Antibiotic resistance by enzymatic modification of antibiotic targets. Trends Mol. Med. 26, 768–782.

7. Arnold, B.J., Huang, I.T., and Hanage, W.P. (2022). Horizontal gene transfer and adaptive evolution in bacteria. Nat. Rev. Microbiol. 20, 206–218.

8. Darby, E.M., Trampari, E., Siasat, P., Gaya, M.S., Alav, I., Webber, M.A., and Blair, J.M.A. (2023). Molecular mechanisms of antibiotic resistance revisited. Nat. Rev. Microbiol. 21, 280–295.

9. Wright, G.D. (2005). Bacterial resistance to antibiotics: enzymatic degradation and modification. Adv. Drug Deliv. Rev. 57, 1451–1470.

10. Tooke, C.L., Hinchliffe, P., Bragginton, E.C., Colenso, C.K., Hirvonen, V.H.A., Takebayashi, Y., and Spencer, J. (2019). *β*-Lactamases and *β*-lactamase inhibitors in the 21st century. J. Mol. Biol. 431, 3472–3500.

11. Maher, C., and Hassan, K.A. (2023). The Gram-negative permeability barrier: tipping the balance of the in and the out. mBio 14, e0120523.

12. Xu, Y., Wei, W., Lei, S., Lin, J., Srinivas, S., and Feng, Y. (2018). An evolutionarily conserved mechanism for intrinsic and transferable polymyxin resistance. mBio 9, e02317–17.

13. Forde, B.M., Zowawi, H.M., Harris, P.N.A., Roberts, L., Ibrahim, E., Shaikh, N., Deshmukh, A., Sid Ahmed, M.A., Al Maslamani, M., Cottrell, K., et al. (2018). Discovery of *mcr-1*-mediated colistin resistance in a highly virulent *Escherichia coli* lineage. mSphere 3, e00486–18.

14. Shahzad, S., Willcox, M.D.P., and Rayamajhee, B. (2023). A review of resistance to polymyxins and evolving mobile colistin resistance gene (mcr) among pathogens of clinical significance. Antibiotics 12, 1597.

15. Mondal, A.H., Khare, K., Saxena, P., Debnath, P., Mukhopadhyay, K., and Yadav, D. (2024). A review on colistin resistance: an antibiotic of last resort. Microorganisms 12, 772.

16. Novović, K., and Jovčić, B. (2023). Colistin resistance in *Acinetobacter baumannii*: molecular mechanisms and epidemiology. Antibiotics 12, 516.

17. Olaitan, A.O., Morand, S., and Rolain, J.M. (2014). Mechanisms of polymyxin resistance: acquired and intrinsic resistance in bacteria. Front. Microbiol. 5, 643.

18. Mohapatra, S.S., Dwibedy, S.K., and Padhy, I. (2021). Polymyxins, the last-resort antibiotics: mode of action, resistance emergence, and potential solutions. J. Biosci. 46, 85.

19. Gogry, F.A., Siddiqui, M.T., Sultan, I., and Haq, Q.M.R. (2021). Current update on intrinsic and acquired colistin resistance mechanisms in bacteria. Front. Med. 8, 677720.

20. Aghapour, Z., Gholizadeh, P., Ganbarov, K., Bialvaei, A.Z., Mahmood, S.S., Tanomand, A., Yousefi, M., Asgharzadeh, M., Yousefi, B., and Kafil, H.S. (2019). Molecular mechanisms related to colistin resistance in Enterobacteriaceae. Infect. Drug Resist. 12, 965–975.

21. Suardíaz, R., Lythell, E., Hinchliffe, P., van der Kamp, M., Spencer, J., Fey, N., and Mulholland, A.J. (2021). Catalytic mechanism of the colistin resistance protein MCR-1. Org. Biomol. Chem. 19, 3813–3819.

22. Mmatli, M., Mbelle, N.M., and Osei Sekyere, J. (2025). Plasmid-borne *mcr-1* and replicative transposition of episomal and chromosomal *bla*NDM-1, *bla*OXA-69, and *bla*OXA-23 carbapenemases in a clinical *Acinetobacter baumannii* isolate. mSystems 10, e0168324.

23. Dhanda, G., Acharya, Y., and Haldar, J. (2023). Antibiotic adjuvants: a versatile approach to combat antibiotic resistance. ACS Omega 8, 10757–10783.

24. Joseph, J., Boby, S., Mooyottu, S., and Muyyarikkandy, M.S. (2025). Antibiotic potentiators as a promising strategy for combating antibiotic resistance. NPJ Antimicrob. Resist. 3, 53.

25. Sargianou, M., Stathopoulos, P., Vrysis, C., Tzvetanova, I.D., and Falagas, M.E. (2025). New *β*-lactam/*β*-lactamase inhibitor combination antibiotics. Pathogens 14, 307.

26. MacNair, C.R., Stokes, J.M., Carfrae, L.A., Fiebig-Comyn, A.A., Coombes, B.K., Mulvey, M.R., and Brown, E.D. (2018). Overcoming *mcr-1* mediated colistin resistance with colistin in combination with other antibiotics. Nat. Commun. 9, 458.

27. Lan, X.J., Yan, H.T., Lin, F., Hou, S., Li, C.C., Wang, G.S., Sun, W., Xiao, J.H., and Li, S. (2019). Design, synthesis and biological evaluation of 1-phenyl-2-(phenylamino) ethanone derivatives as novel MCR-1 inhibitors. Molecules 24, 2719.

28. Tang, F., Peng, W., Kou, X., Chen, Z., and Zhang, L. (2024). High-throughput screening identification of apigenin that reverses the colistin resistance of *mcr-1*-positive pathogens. Microbiol. Spectr. 12, e00341–24.

29. Hanpaibool, C., Ngamwongsatit, N., Ounjai, P., Yotphan, S., Wolschann, P., Mulholland, A.J., Spencer, J., and Rungrotmongkol, T. (2023). Pyrazolones potentiate colistin activity against MCR-1-producing resistant bacteria: computational and microbiological study. ACS Omega 8, 8366–8376.

30. Guo, W., Liu, Y., Yao, Z., Zhou, H., Wang, X., Huang, Z., Zhang, X., Wu, Q., and Zhou, T. (2023). Bithionol restores sensitivity of multidrug-resistant gram-negative bacteria to colistin with antimicrobial and anti-biofilm effects. ACS Infect. Dis. 9, 1634–1646.

31. Liang, L., Zhong, L.L., Wang, L., Zhou, D., Li, Y., Li, J., Chen, Y., Liang, W., Wei, W., Zhang, C., et al. (2023). A new variant of the colistin resistance gene MCR-1 with co-resistance to *β*-lactam antibiotics reveals a potential novel antimicrobial peptide. PLoS Biol. 21, e3002433.

32. Talevi, A. (2024). Computer-aided drug discovery and design: recent advances and future prospects. Methods Mol. Biol. 2714, 1–20.

33. Zhang, J.Z.H. (2026). Recent advances in computer-aided drug design and drug discovery. Molecules 31, 1606.

34. Tingle, B.I., Tang, K.G., Castanon, M., Gutierrez, J.J., Khurelbaatar, M., Dandarchuluun, C., Moroz, Y.S., and Irwin, J.J. (2023). ZINC-22—A free multi-billion-scale database of tangible compounds for ligand discovery. J. Chem. Inf. Model. 63, 1166–1176.

35. Aron, Z.D., Mehrani, A., Hoffer, E.D., Connolly, K.L., Srinivas, P., Torhan, M.C., Alumasa, J.N., Cabrera, M., Hosangadi, D., Barbor, J.S., Cardinale, S.C., Kwasny, S.M., Morin, L.R., Butler, M.M., Opperman, T.J., Bowlin, T.L., Jerse, A., Stagg, S.M., Dunham, C.M., and Keiler, K.C. (2021). *trans*-Translation inhibitors bind to a novel site on the ribosome and clear *Neisseria gonorrhoeae* in vivo. Nat. Commun. 12, 1799.

36. Barber, T.S., and Alumasa, J.N. (2026). Antibiotic-adjuvants abolish resistance conferred by the *Staphylococcus aureus* erythromycin resistance methyltransferase in an *Escherichia coli* model. ACS Infect. Dis. 12, 376–389.

37. Zinkle, A.P., Batista, M.B., Herrera, C.M., Erramilli, S.K., Kloss, B., Ashraf, K.U., Nosol, K., Zhang, G., Cater, R.J., Marty, M.T., et al. (2025). Mechanistic basis of antimicrobial resistance mediated by the phosphoethanolamine transferase MCR-1. Nat. Commun. 16, 10516.

38. Irwin, J.J., Tang, K.G., Young, J., Dandarchuluun, C., Wong, B.R., Khurelbaatar, M., Moroz, Y.S., Mayfield, J., and Sayle, R.A. (2020). ZINC20—A free ultralarge-scale chemical database for ligand discovery. J. Chem. Inf. Model. 60, 6065–6073.

39. Daina, A., Michielin, O., and Zoete, V. (2017). SwissADME: a free web tool to evaluate pharmacokinetics, drug-likeness and medicinal chemistry friendliness of small molecules. Sci. Rep. 7, 42717.

40. Keiser, M.J., Roth, B.L., Armbruster, B.N., Ernsberger, P., Irwin, J.J., and Shoichet, B.K. (2007). Relating protein pharmacology by ligand chemistry. Nat. Biotechnol. 25, 197–206.

41. Crowley, L.C., Scott, A.P., Marfell, B.J., Boughaba, J.A., Chojnowski, G., and Waterhouse, N.J. (2016). Measuring cell death by propidium iodide uptake and flow cytometry. Cold Spring Harb. Protoc. 2016, pdb.prot087163.

42. Andrade, F.F., Silva, D., Rodrigues, A., and Pina-Vaz, C. (2020). Colistin update on its mechanism of action and resistance, present and future challenges. Microorganisms 8, 1716.

43. Sabnis, A., Hagart, K.L., Klöckner, A., Becce, M., Evans, L.E., Furniss, R.C.D., Mavridou, D.A., Murphy, R., Stevens, M.M., Davies, J.C., et al. (2021). Colistin kills bacteria by targeting lipopolysaccharide in the cytoplasmic membrane. eLife 10, e65836.

44. Kern, W.V., Steinke, P., Schumacher, A., Schuster, S., von Baum, H., and Bohnert, J.A. (2006). Effect of 1-(1-naphthylmethyl)-piperazine, a novel putative efflux pump inhibitor, on antimicrobial drug susceptibility in clinical isolates of *Escherichia coli*. J. Antimicrob. Chemother. 57, 339–343.

45. Zhang, L., Tian, X., Sun, L., Mi, K., Wang, R., Gong, F., and Huang, L. (2024). Bacterial efflux pump inhibitors reduce antibiotic resistance. Pharmaceutics 16, 170.

46. Jerabek-Willemsen, M., Wienken, C.J., Braun, D., Baaske, P., and Duhr, S. (2011). Molecular interaction studies using microscale thermophoresis. Assay Drug Dev. Technol. 9, 342–353.

47. Marimuthu, S.K., Ramakrishnan, V., and Thamotharan, S. (2025). Targeting dual substrate pockets of colistin resistance conferring MCR-1 of *Escherichia coli* with natural products: insights from high throughput virtual screening and molecular dynamics simulations. J. Biomol. Struct. Dyn. 43, 7139–7154.

48. Sun, Z., and Palzkill, T. (2021). Deep Mutational Scanning Reveals the Active-Site Sequence Requirements for the Colistin Antibiotic Resistance Enzyme MCR-1. mBio 12, e02776–21.

49. Lenth, R., and Piaskowski, J. (2026). emmeans: Estimated Marginal Means, aka Least-Squares Means. R package version 2.0.5. https://rvlenth.github.io/emmeans/.

50. Wickham, H., Averick, M., Bryan, J., Chang, W., D’Agostino McGowan, L., François, R., Grolemund, G., Hayes, A., Henry, L., Hester, J., et al. (2019). Welcome to the Tidyverse. J. Open Source Softw. 4, 1686.

