## Supplemental Figures for "Isoindole-benzohydrazide derivative potentiates colistin activity to overcome resistance conferred by MCR-1 variants in *Escherichia coli*"

for

### SUPPORTING INFORMATION: FIGURES

#### Figure S1: Compound structures, identifiers, and predicted binding energies

**A**

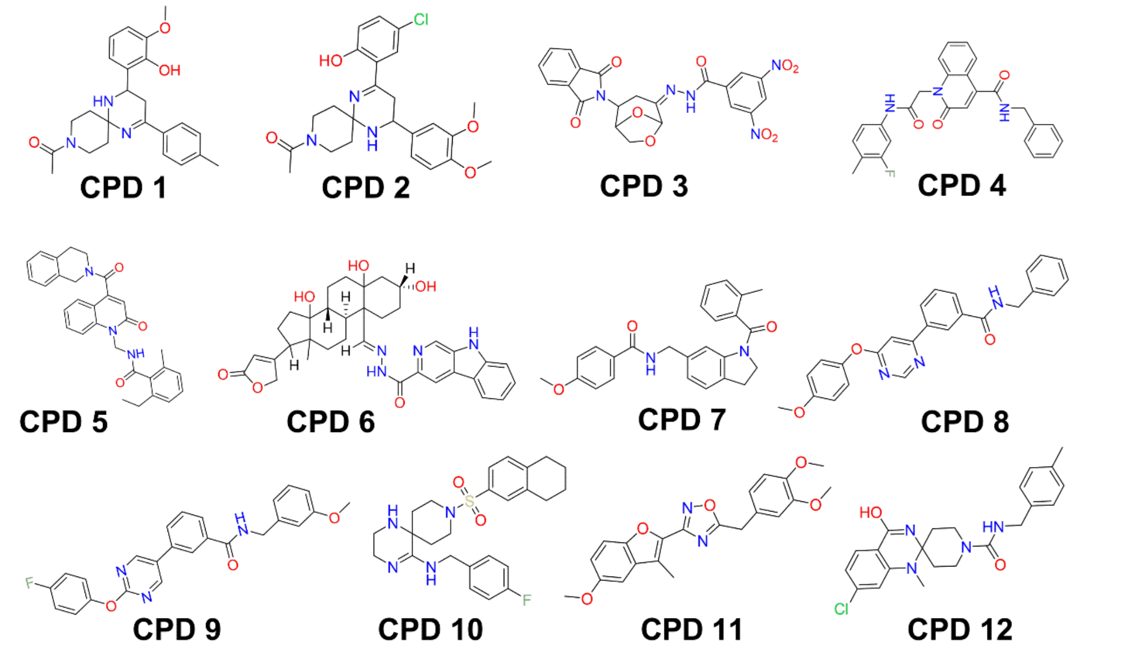


**B**

<table>
<tr><th>Compound ID</th><th>Binding Energy (kcal/mol)</th><th>SMILES</th></tr>
<tr><td>CDP-1</td><td>-8.4</td><td>COc1cccc(C2CC(=NC3(CCN(CC3)C(C)=O)N2)c2ccc(C)cc2)c1O |c:9|</td></tr>
<tr><td>CPD-2</td><td>-8.7</td><td>COc1ccc(cc1OC)C1CC(=NC2(CCN(CC2)C(C)=O)N1)c1cc(Cl)ccc1O |c:13|</td></tr>
<tr><td>CPD-3</td><td>-9.7</td><td>[O-][N+](=O)c1cc(cc(c1)[N+]([O-])=O)C(=O)N\N=C1\CC(C2COC1O2)N1C(=O)c2ccccc2C1=O</td></tr>
<tr><td>CPD-4</td><td>-9.5</td><td>Cc1ccc(NC(=O)Cn2c3ccccc3c(cc2=O)C(=O)NCc2ccccc2)cc1F</td></tr>
<tr><td>CPD-5</td><td>-9.2</td><td>CCc1cccc(C)c1NC(=O)Cn1c2ccccc2c(cc1=O)C(=O)N1CCc2ccccc2C1</td></tr>
<tr><td>CPD-6</td><td>-9.2</td><td>CC12CC[C@H]3[C@@H](CCC4(O)C[C@H](O)CCC34\C=N\NC(=O)c3cc4c(cn3)[nH]c3ccccc43)C1(O)CCC2C1=CC(=O)OC1 |r,t:46|</td></tr>
<tr><td>CPD-7</td><td>-8.1</td><td>COc1ccc(cc1)C(=O)NCc1ccc2CCN(C(=O)c3ccccc3C)c2c1</td></tr>
<tr><td>CPD-8</td><td>-8.2</td><td>COc1ccc(Oc2cc(ncn2)-c2cccc(c2)C(=O)NCc2ccccc2)cc1</td></tr>
<tr><td>CPD-9</td><td>-9.1</td><td>COc1cccc(CNC(=O)c2cccc(c2)-c2cnc(Oc3ccc(F)cc3)nc2)c1</td></tr>
<tr><td>CPD-10</td><td>-8.8</td><td>Cl.Fc1ccc(CNC2=NCCNC22CCN(CC2)S(=O)(=O)c2ccc3CCCCc3c2)cc1 |t:7|</td></tr>
<tr><td>CPD-11</td><td>-7.6</td><td>COc1ccc2oc(c(C)c2c1)-c1noc(Cc2ccc(OC)c(OC)c2)n1</td></tr>
<tr><td>CPD-12</td><td>-9.3</td><td>CN1c2cc(Cl)ccc2C(=O)NC11CCN(CC1)C(=O)NCc1ccc(C)cc1</td></tr>
</table>

**Figure S1: Structures of test compounds and their predicted binding energies to MCR-1. A)** Structures of the 12 test compounds prioritized from the in silico computational screens conducted to identify compounds with ideal drug-like properties, predicted to bind MCR-1. **B)** Results from the in silico virtual screens using AutoDock Vina demonstrating the predicted binding energies for the 12 compounds prioritized in this study and their identifiers. Abbreviation: CPD – compound.

**Figure S2: Colistin-JNAL-042 combinatorial checkerboard assays**

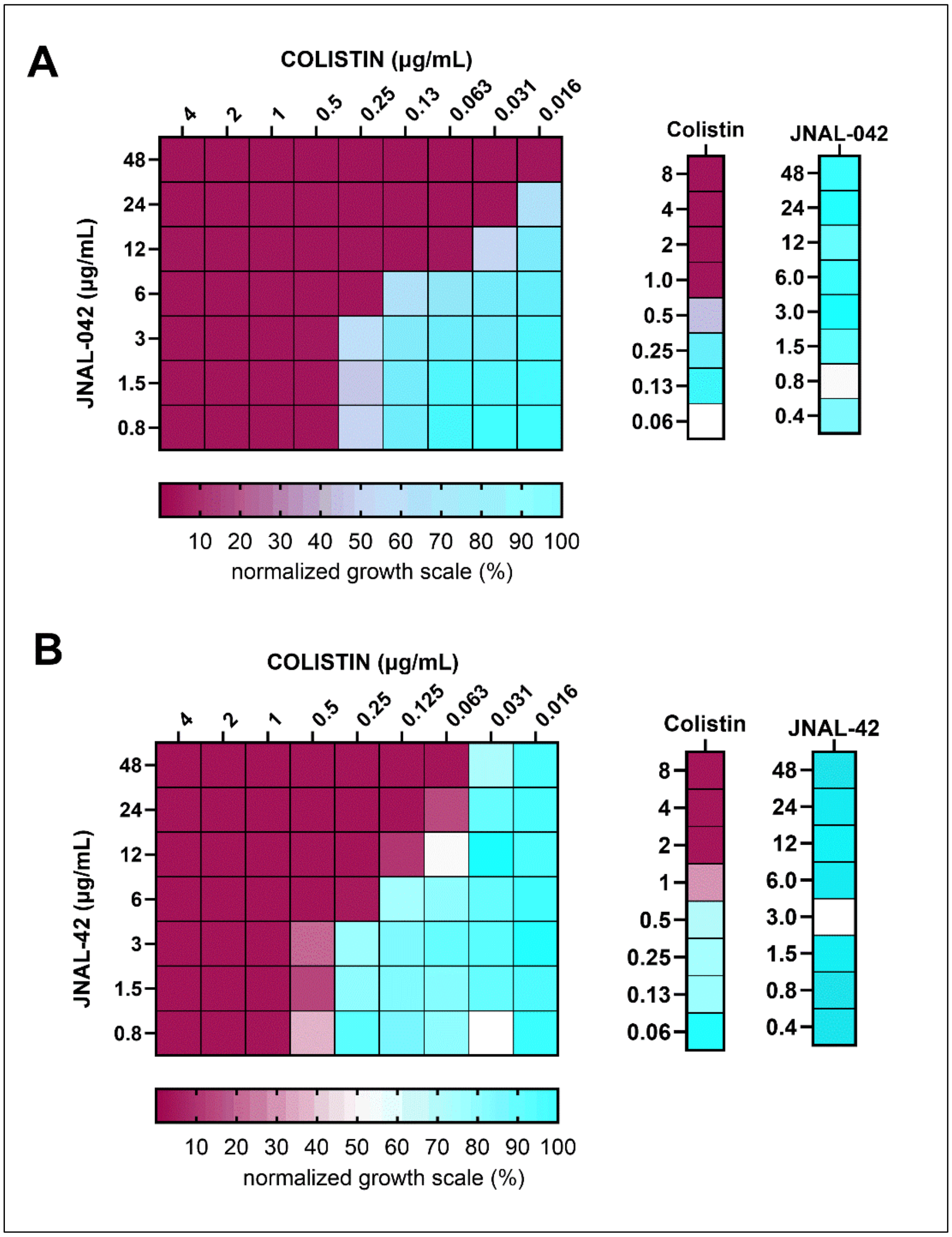


**Figure S2. JNAL-042 lacks intrinsic antibiotic activity but potentiates that of colistin against MCR-M6-mediated resistance.** Checkerboard assays evaluating synergy between colistin and JNAL-042 using strain ASW-018 expressing the MCR-M6 variant. The figure shows heatmaps from two biological replicates evaluating colistin independently from 0 – 8 µg/mL and JNAL-042 from 0 – 48 µg/mL. For the combinatorial section, the adjuvant is serially diluted down the plate, and colistin is diluted across the plate. The data were normalized to drug-free control wells to obtain the corresponding growth scale. The MIC for colistin was in the 1–2 µg/mL range, while JNAL-042 showed no appreciable activity up to 48 µg/mL.

#### Figure S3: Effect of JNAL-042 on colistin activity in time-kill kinetic assays

##### Bio-rep_2

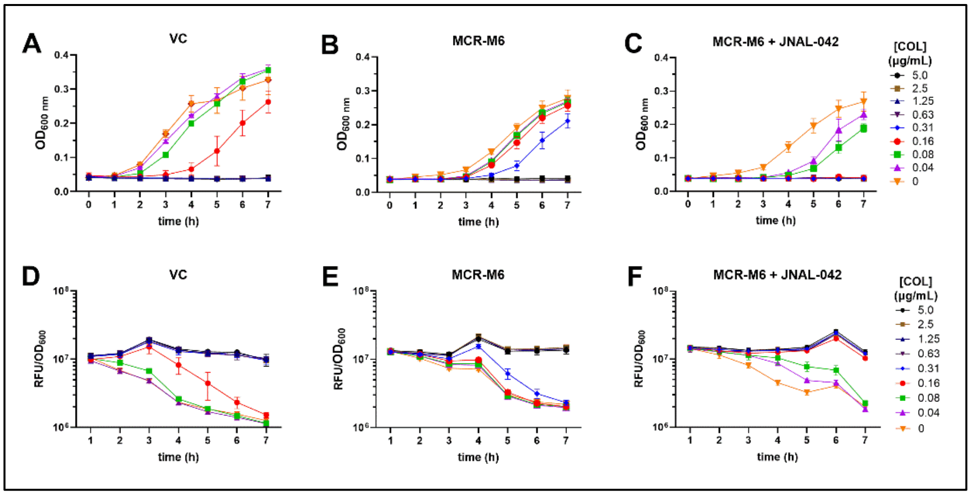


##### Bio-rep_3

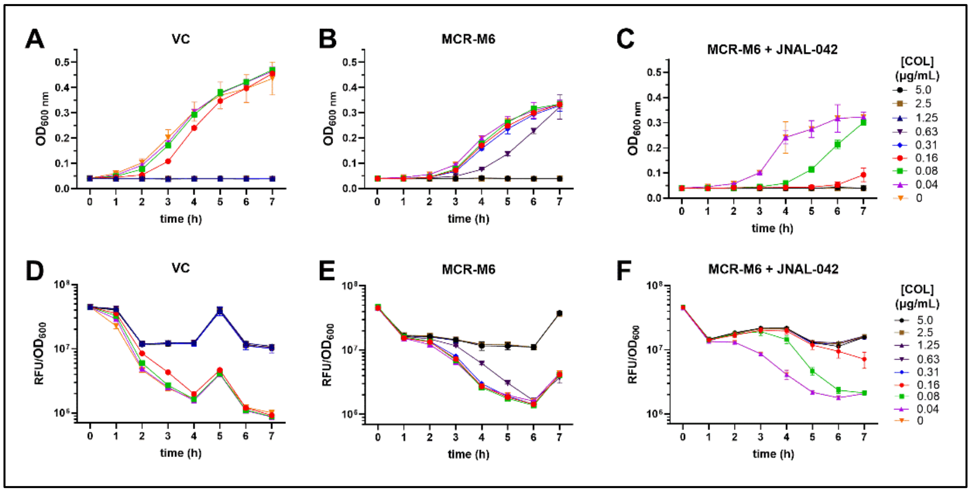


**Figure S3. JNAL-042 potentiates COL activity to abolish MCR-1-mediated resistance in time-kill assays with efflux-deficient *E. coli* strains**. The figure presents data from 2 additional biological replicates of experiments used to generate **Figure 3** in the main manuscript. The experiment employed an engineered empty vector-carrying *E. coli* strain (VC; ASW-021; a COL-sensitive strain; A, D, & G) and one containing the gene encoding the MCR-M6 variant (a COL-resistant strain; B-C, E-F, & H-I). **A-C**) Time-dependent growth profiles of the VC and MCR-M6-expressing strains treated with varying COL concentrations monitored using cell density at $OD_{600nm}$. **D-F**) Time-kill assay monitored using the PI incorporation assay, demonstrating the restoration of COL activity in the presence of JNAL-042 (12 μg/mL) to an effective bactericidal concentration (EBC) similar to that observed in the VC strain (0.63 μg/mL). For graphs A−F in each biological replicate, the data presented are averages from three technical replicates (±SD). Abbreviations: VC – Vector Control; OD – Optical Density; RFU – Relative Fluorescence Units; CFU – Colony-Forming Units.

#### Figure S4: Growth & PI Time-kill Assays for Polymyxin B (PMB)

##### Bio-rep_1

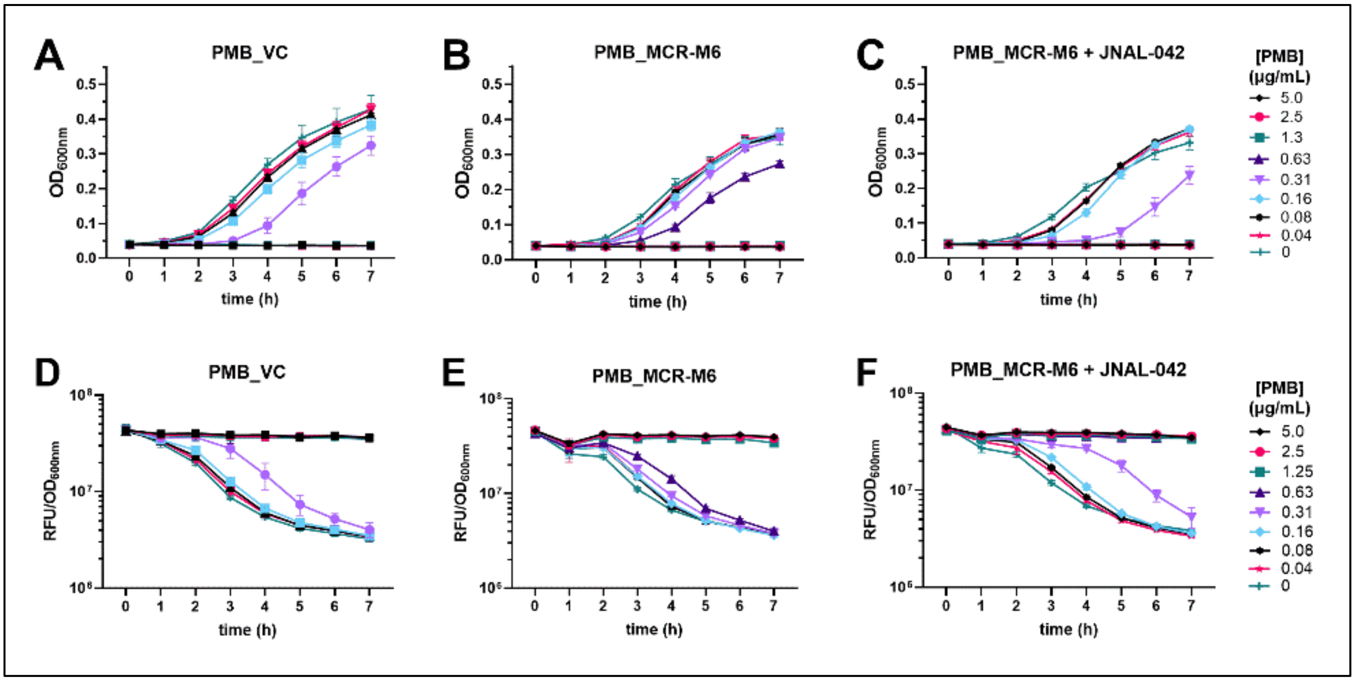


##### Bio-rep_2

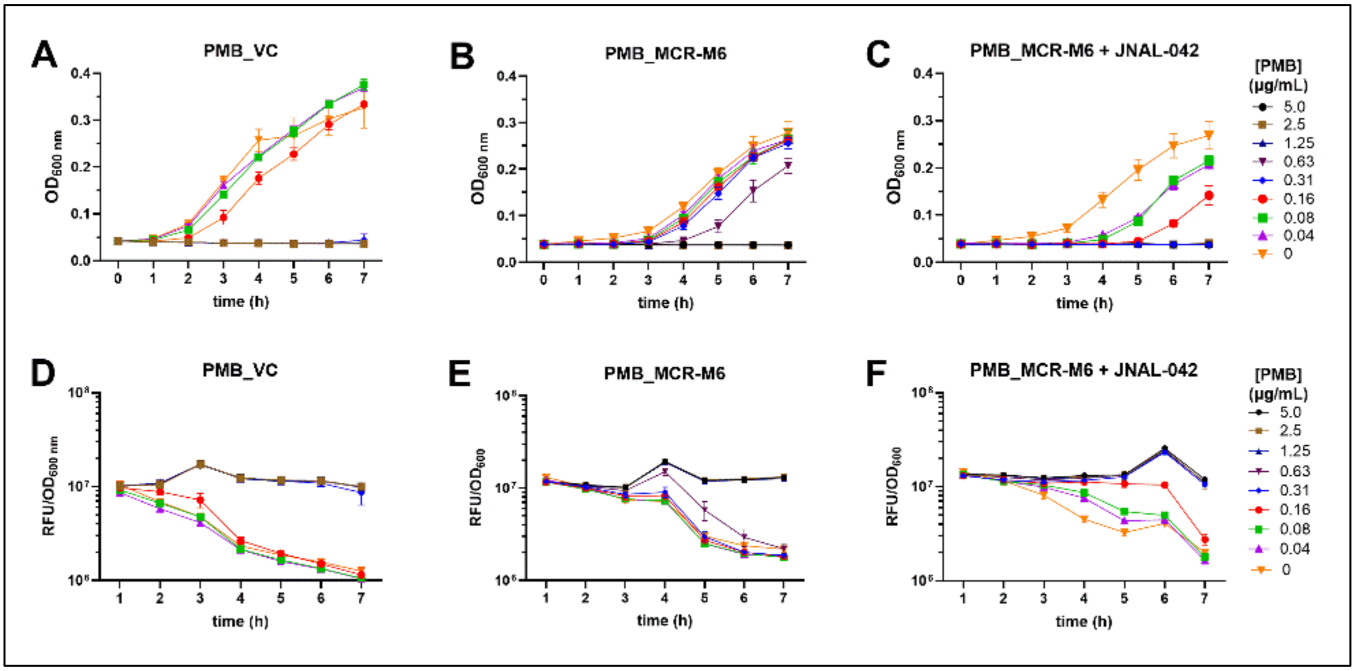

#### Bio-rep_3

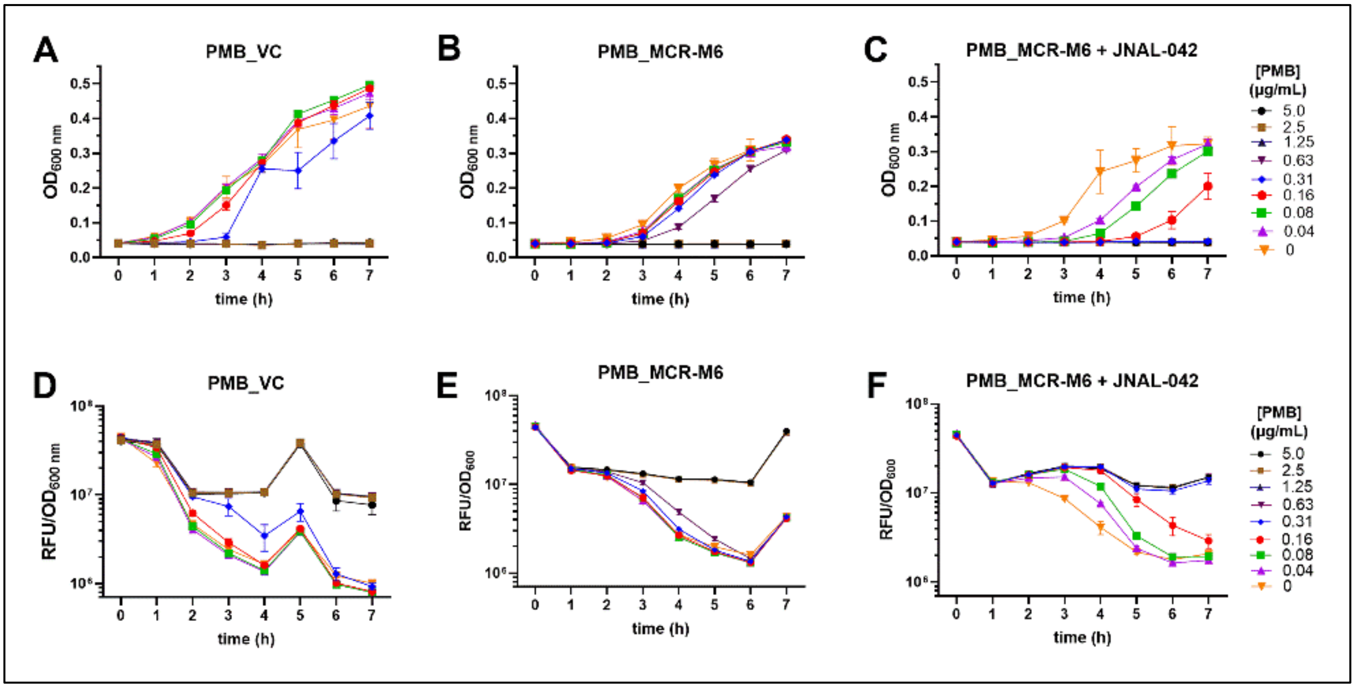


**Figure S4. JNAL-042 potentiates polymyxin B (PMB) activity to abolish MCR-1-mediated resistance in time-kill assays using efflux-deficient *E. coli* strains**. The figure presents data from 3 biological replicates of PMB experiments conducted in parallel with those evaluating colistin used to generate **Figure 3** in the main manuscript. The experiment employed an engineered empty vector-carrying *E. coli* strain (VC; ASW-021; a PMB-sensitive strain; A, D, & G) and one containing the gene encoding the MCR-M6 variant (a PMB-resistant strain; B-C, E-F, & H-I). **A-C**) Time-dependent growth profiles of the VC and MCR-M6-expressing strains treated with varying PMB concentrations monitored using cell density at $OD_{600nm}$. **D-F**) Results from time-kill assays monitored using the PI incorporation assay, demonstrating the restoration of PMB activity in the presence of JNAL-042 (12 µg/mL) to an effective bactericidal concentration (EBC) similar to that observed in the VC strain (0.31 µg/mL). For graphs A−F in each biological replicate, the data presented are averages from three technical replicates (±SD). Abbreviations: VC – vector control; OD – Optical Density; RFU – Relative Fluorescence Units; CFU – Colony-Forming Units.

**Figure S5 – Post-induction/delayed-treatment time-kill assay biological replicates**

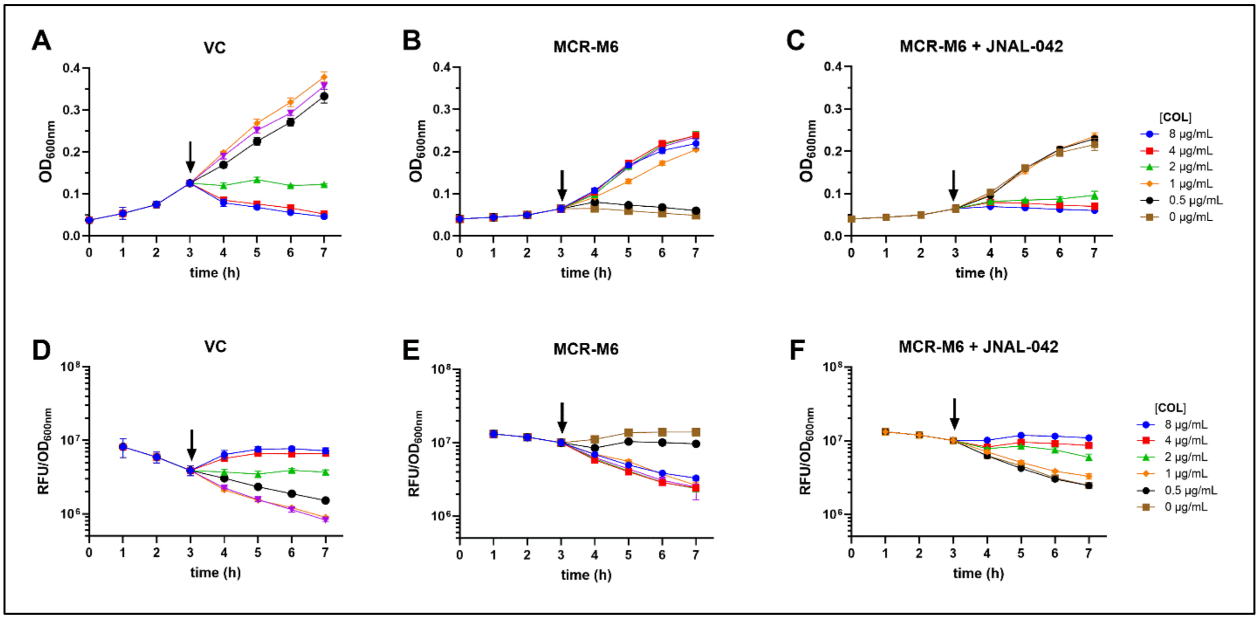


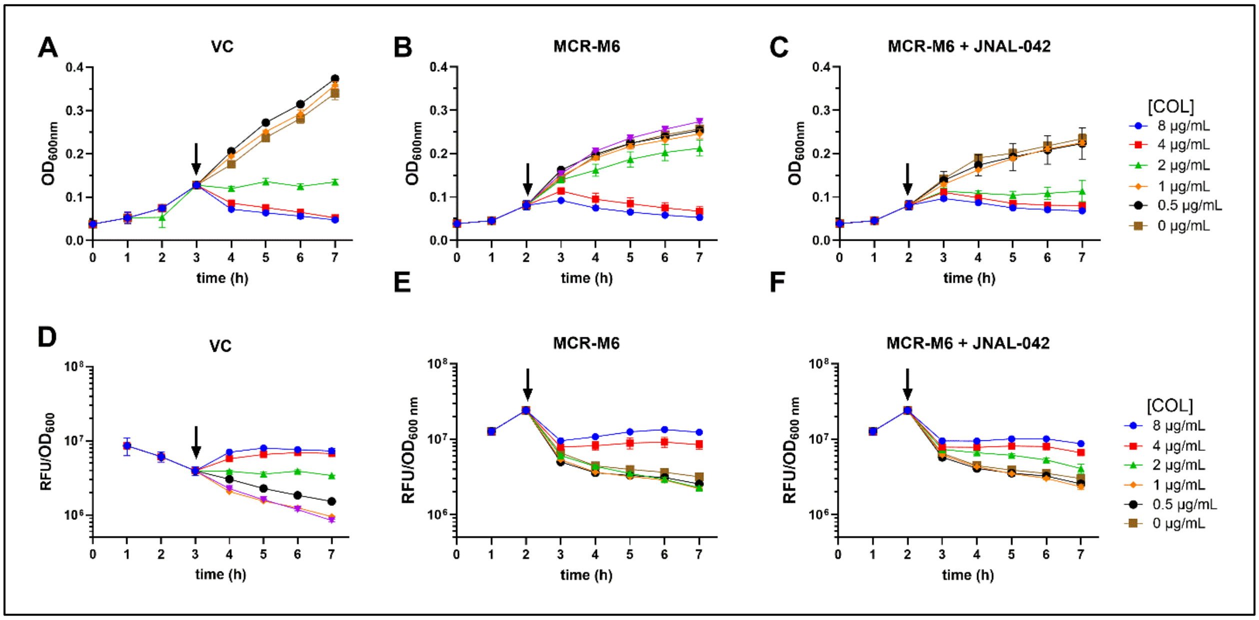


**Figure S5. JNAL-042 potentiates COL activity against MCR-M6-mediated resistance in post-induction delayed-treatment kinetic bacterial susceptibility assays**. The figure presents data from 2 additional biological replicates of experiments used to generate **Figure 4** in the main manuscript. **A-C**) Growth profiles monitored using cell density ($OD_{600nm}$) for control (ASW-021) and MCR-M6-expressing strain (ASW-018), evaluating the effect of adding COL (A, B) or a combination of COL + 24 µg/mL JNAL-042 (C) 3 h (arrow) after inducing enzyme production. **D-F**) Time-dependent bacterial cell death profiles monitored over 7 h using the PI incorporation assays for the corresponding cultures in A-C. The data for each biological replicate are averages from three technical replicates (±SD). Abbreviations: VC – vector control; OD – Optical Density; RFU – Relative Fluorescence Units; CFU – Colony-Forming Units.

**Figure S6: Antibacterial susceptibility screens with 1-(1-naphthylmethyl)piperazine (NMP)**

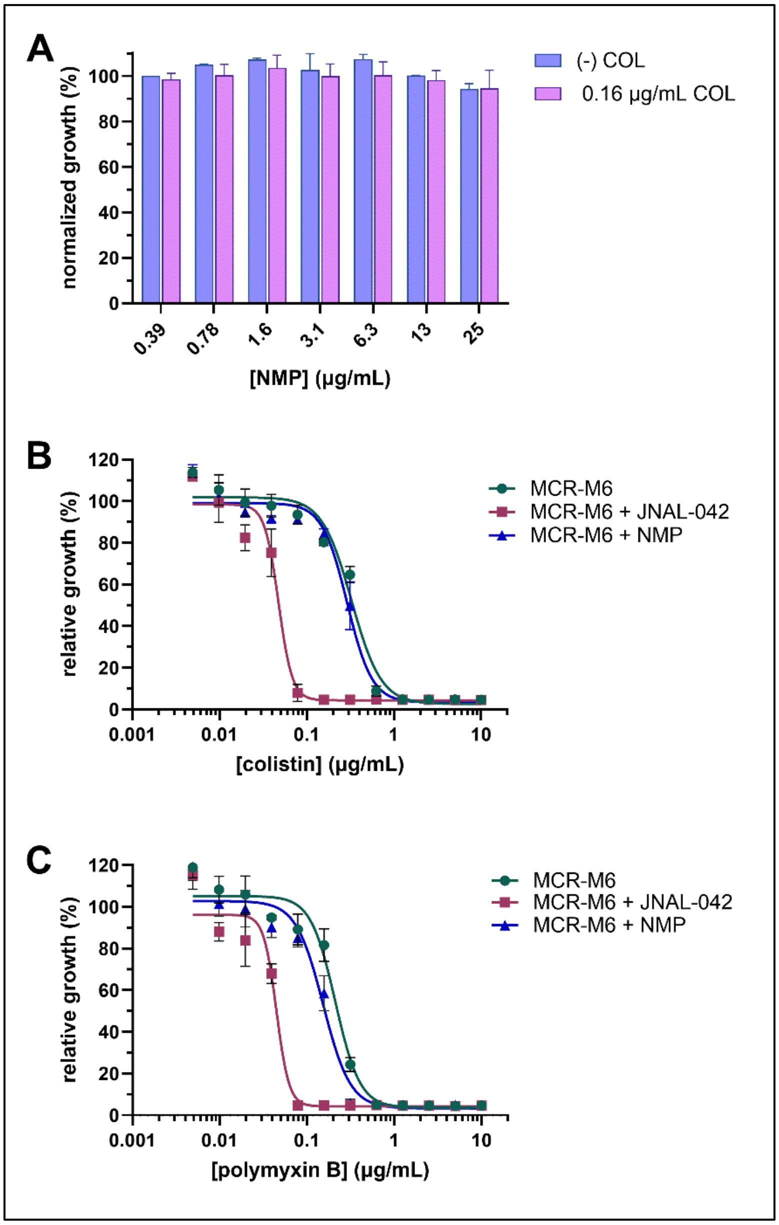


**Figure S6. 1-(1-naphthylmethyl)piperazine (NMP) lacks antibacterial growth-inhibitory properties**. Experiments evaluating potential antibacterial properties of NMP at concentrations used in the assay. **A**) Growth profiles of strain ASW-020 with or without COL, evaluating potential arrest by NMP, demonstrating no appreciable activity at all the concentrations evaluated. **B, C**) Results from antibacterial susceptibility assays with strain ASW-018 highlighting the dose-response profiles for COL or PMB alone (circles), combined with JNAL-042 (12 µg/mL; squares) or NMP (17 µg/mL; triangles). The data show no appreciable impact of NMP at 17 µg/mL on colistin activity; this concentration was selected for use in the time-kill assays against the WT *E. coli* strains.

#### Figure S7: Combinatorial time-kill assays involving COL, PMB, JNAL-042, and NMP

##### Biological replicate #2

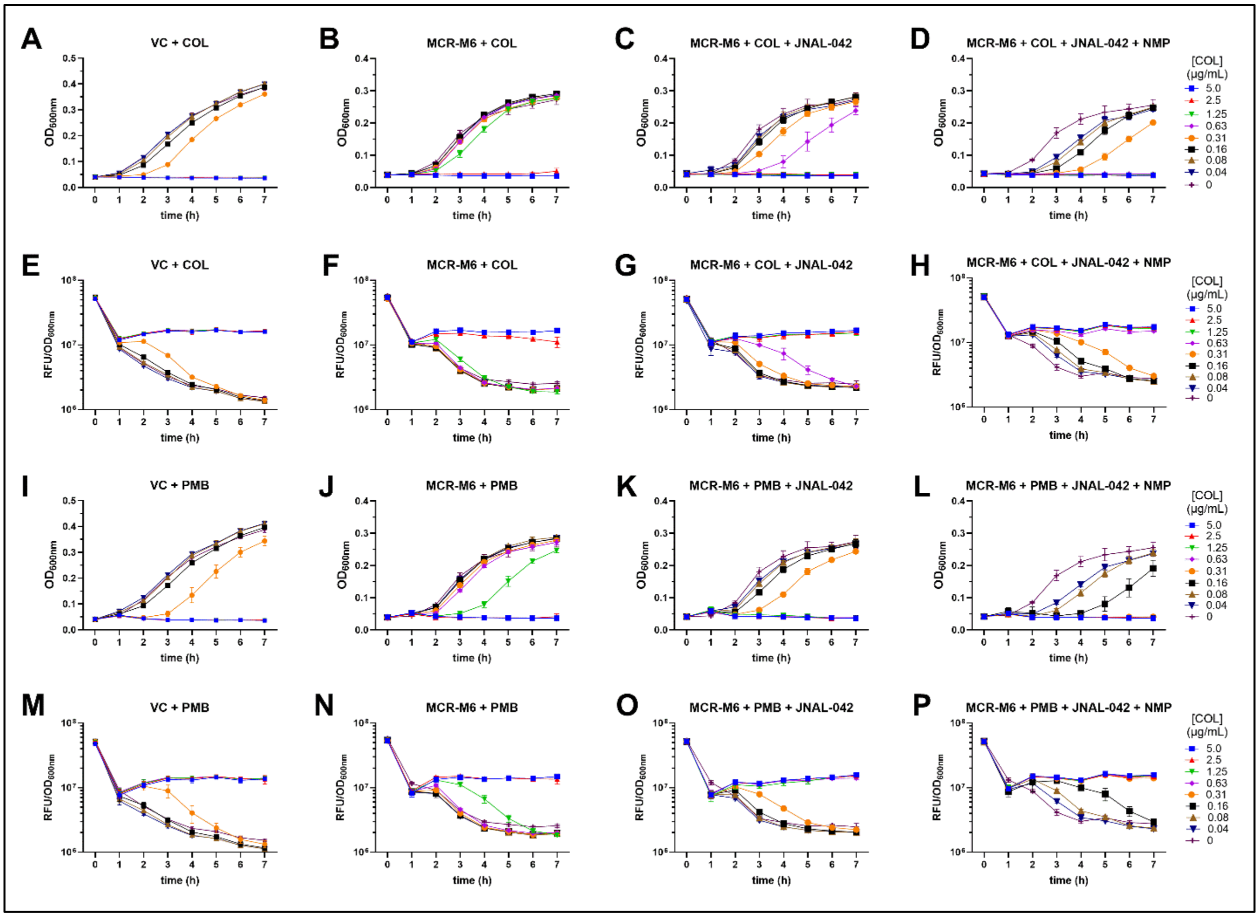


**Figure S7. JNAL-042 suppresses MCR-M6-mediated resistance in WT *E. coli* cells**. The figure presents data from biological replicate #2 of experiments used to generate **Figure 5** in the main manuscript. Results from experiments assessing COL activity in actively growing cultures by monitoring time-dependent growth ($OD_{600nm}$) (A-D) and cell death using a fluorescence-based PI incorporation reporter assay (E-H) in WT *E. coli* strains carrying an empty vector (ASW-022) or an MCR-M6-expression plasmid (ASW-020). **A, B, E, F**) Results obtained from monitoring COL activity. **C, G**) Activity profiles of COL-JNAL-042 (24 µg/ml) combination treatments in the presence of the enzyme. **D, H**) Data highlighting the activity of a treatment cocktail, including COL + JNAL-042 (24 µg/ml) + NMP (17 µg/ml), demonstrating complete restoration of COL activity. **I, J, M, N**) Results obtained from monitoring PMB activity. **K, O**) Activity profiles of PMB-JNAL-042 (24 µg/ml) combination treatments in the presence of the MCR-M6. **L, P**) Results demonstrating the activity of a treatment cocktail, including COL + JNAL-042 (24 µg/ml) + NMP (17 µg/ml), demonstrating complete restoration of COL activity. The data are averages from three technical replicates (±SD). Abbreviations: VC: vector control; RFU: relative fluorescence units; NMP: 1-(1-naphthylmethyl)piperazine.

**Figure S8: Combinatorial time-kill assays involving COL, PMB, JNAL-042, and NMP**

**Biological replicate #3**

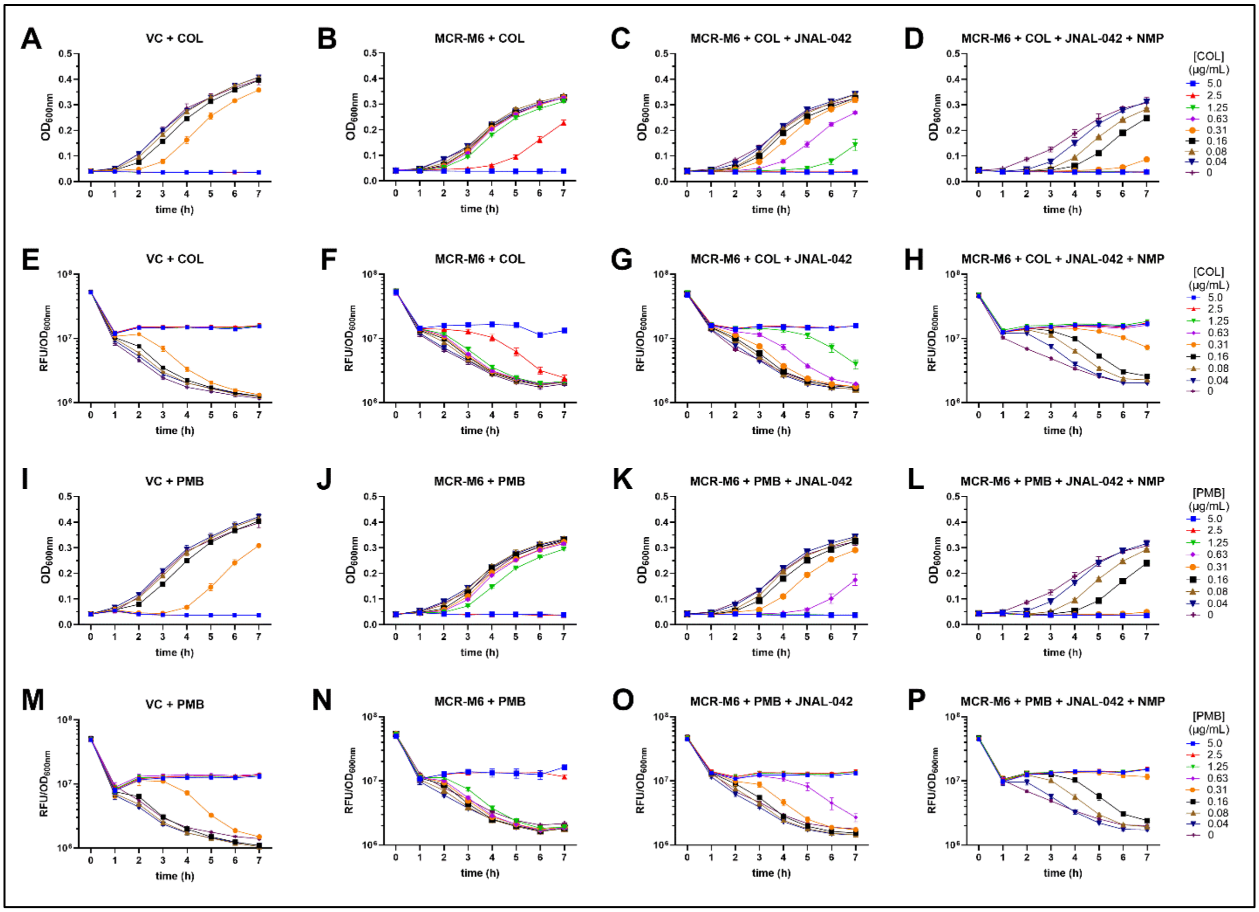


**JNAL-042 suppresses MCR-M6-mediated resistance in WT *E. coli* cells**. The figure presents data from biological replicate #3 of experiments used to generate **Figure 5** in the main manuscript. Results from experiments assessing COL activity in actively growing cultures by monitoring time-dependent growth (OD$_{600nm}$) (A-D) and cell death using a fluorescence-based PI incorporation reporter assay (E-H) in WT *E. coli* strains carrying an empty vector (ASW-022) or an MCR-M6-expression plasmid (ASW-020). **A, B, E, F**) Results obtained from monitoring COL activity. **C, G**) Activity profiles of COL-JNAL-042 (24 µg/ml) combination treatments in the presence of the enzyme. **D, H**) Data highlighting the activity of a treatment cocktail, including COL + JNAL-042 (24 µg/ml) + NMP (17 µg/ml), demonstrating complete restoration of COL activity. **I, J, M, N**) Results obtained from monitoring PMB activity. **K, O**) Activity profiles of PMB-JNAL-042 (24 µg/ml) combination treatments in the presence of the MCR-M6. **L, P**) Results demonstrating the activity of a treatment cocktail, including COL + JNAL-042 (24 µg/ml) + NMP (17 µg/ml), demonstrating complete restoration of COL activity. The data are averages from three technical replicates (±SD). Abbreviations: VC: vector control; RFU: relative fluorescence units; NMP: 1-(1-naphthylmethyl)piperazine

**Figure S9: JNAL-042 docking and structural alignment of MCR variants**

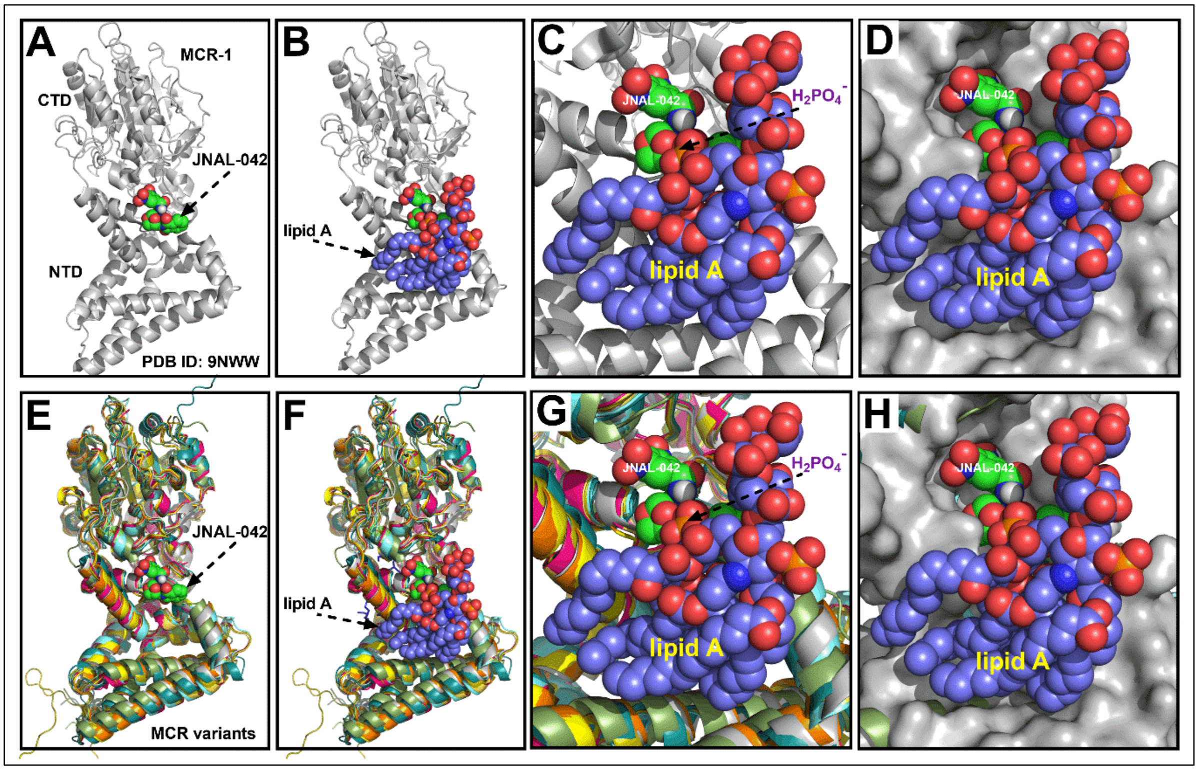


**Figure S9. JNAL-042 may possess broad-spectrum activity against MCR variants. A**) Cartoon representation of the cryo-EM structure of MCR-1 (PDB ID: 9NWW; lipid A and PEtN have been removed for clarity) demonstrating one docked conformation of JNAL-042. **B**) The same structure of JNAL-042 bound to MCR-1, highlighting the predicted binding site of the adjuvant relative to that of lipid A. C) Zoomed-in view of the predicted binding site of JNAL-042, highlighting its proximity to a phosphate functional group, typically modified by MCR-1 in the second catalytic step. **D**) Same image as C, with the protein represented in the filled surface view. **E**). Cartoon representation of the alignment of MCR-variant (1–10) structures obtained from the AlphaFold database or predicted using the software, showing high structural similarity and a potentially conserved binding site for JNAL-042. **F**) MCR-variant structures overlaid with MCR-1, demonstrating the conserved binding site for lipid A. **G**, **H**) Zoomed-in views of the cartoon and surface representation, respectively, depicting conservation of the adjuvant binding pocket. Abbreviations: CTD – C-terminal domain; NTD – N-terminal domain.
